# Biparatopic sybody constructs neutralize SARS-CoV-2 variants of concern and mitigate emergence of drug resistance

**DOI:** 10.1101/2020.11.10.376822

**Authors:** Justin D. Walter, Cedric A.J. Hutter, Alisa A. Garaeva, Melanie Scherer, Iwan Zimmermann, Marianne Wyss, Jan Rheinberger, Yelena Ruedin, Jennifer C. Earp, Pascal Egloff, Michèle Sorgenfrei, Lea M. Hürlimann, Imre Gonda, Gianmarco Meier, Sille Remm, Sujani Thavarasah, Gerrit van Geest, Rémy Bruggman, Gert Zimmer, Dirk J. Slotboom, Cristina Paulino, Philippe Plattet, Markus A. Seeger

## Abstract

The ongoing COVID-19 pandemic represents an unprecedented global health crisis. Here, we report the identification of a synthetic nanobody (sybody) pair (Sb#15 and Sb#68) that can bind simultaneously to the SARS-CoV-2 spike-RBD and efficiently neutralize pseudotyped and live-viruses by interfering with ACE2 interaction. Two spatially-discrete epitopes identified by cryo-EM translated into the rational design of bispecific and tri-bispecific fusions constructs, exhibiting up to 100- and 1000-fold increase in neutralization potency. Cryo-EM of the sybody-spike complex further revealed a novel *up-out* RBD conformation. While resistant viruses emerged rapidly in the presence of single binders, no escape variants were observed in presence of the bispecific sybody. The multivalent bispecific constructs further increased the neutralization potency against globally-circulating SARS-CoV-2 variants of concern. Our study illustrates the power of multivalency and biparatopic nanobody fusions for the development of clinically relevant therapeutic strategies that mitigate the emergence of new SARS-CoV-2 escape mutants.

## INTRODUCTION

The spike glycoprotein (S-protein) is the most prominent surface-exposed entity of severe acute respiratory syndrome coronavirus 2 (SARS-CoV-2), and possesses the vital molecular machinery required for recognition and fusion with host membranes [1]. To date, most authorized vaccines against coronavirus disease 2019 (COVID-19) rely on exposure of patients solely to the S-protein [2]. Similarly, the S-protein is the exclusive target of currently approved monoclonal antibody therapies for COVID-19 [3]. Unfortunately, recent months have seen the emergence and rapid spread of mutant viral strains conferring amino acid changes in the S-protein which can attenuate neutralization by many convalescent, vaccine-induced, and monoclonal antibodies [4, 5]. Therefore, from a public health perspective, it is imperative to pursue the development of therapeutic strategies that can withstand the continued emergence of SARS-CoV-2 escape mutants.

The S-protein mutations that cause increased virulence and immune evasion are predominantly found in the receptor-binding domain (RBD) [5, 6], which is specifically responsible for host recognition via interaction with human angiotensin-converting enzyme 2 (ACE2) [7]. The RBD harbors two hotspots for antibody recognition. One of these epitopes overlaps with the ACE2 binding interface and is evolutionarily unique in SARS-CoV-2; the second so-called “cryptic” epitope is found in a peripheral region that is conserved among RBDs from several characterized coronaviruses [8]. While individually targeting either epitope with antibodies quickly results in the emergence of escape mutants [9, 10], there is growing evidence that simultaneous engagement of both epitopes via polyvalent antibodies may mitigate viral escape [11, 12].

Here, we present two synthetic single-domain antibodies (sybodies), designated Sb#15 and Sb#68, that recognize non-overlapping epitopes on the RBD. Sybodies offer several advantages over conventional antibodies such as the potential for rapid development, low-cost production in prokaryotic expression systems, and facile engineering [13, 14]. Cryogenic electron microscopy (cryo-EM) revealed that Sb#15 binds within the ACE2 interface, whereas Sb#68 engages the adjacent conserved cryptic epitope. Structural analysis also demonstrated that the dual presence of Sb#15 and Sb#68 resulted in the adoption of a novel RBD conformation that we termed *up/out*. Fusion of Sb#15 and Sb#68 yielded a bispecific construct, termed GS4, that displayed enhanced avidity and neutralization potency relative to the separate sybodies. Exposure of SARS-CoV-2 to the individual sybodies *in vitro* resulted in the rapid emergence of escape mutants, including a Q493R-RBD variant (within the ACE2 epitope) that has recently been observed in COVID-19 patients treated with a monoclonal antibody [15], as well as a novel P384H-RBD mutation in the cryptic epitope. In contrast, no escape mutants were detected upon treatment with GS4. Finally, we found that additional valency engineering via covalent trimerization of GS4, giving a construct we termed Tripod-GS4r, results in further enhancement of viral neutralization potential against the B.1.1.7 (alpha) and B.1.351 (beta) SARS-CoV-2 variants of concern. Overall, our study demonstrates favorable prospects for such multivalent sybodies to be a valuable therapeutic tool vis-à-vis future SARS-CoV-2 variants or comparable forthcoming viral pandemics.

## RESULTS

### Identification of a sybody pair that (i) simultaneously bind to the spike RBD, (ii) compete with ACE2 interaction and (iii) efficiently neutralize viruses

We sought to engineer a pair of synthetic nanobodies (sybodies) which may mitigate viral escape due to the simultaneous binding of discrete non-overlapping epitopes of the SARS-CoV-2 spike (S) glycoprotein. Using our established sybody generation workflow [13, 16], we conducted a selection campaign against the isolated receptor-binding domain (RBD) of the S-protein. Upon screening single sybody clones with ELISA and grating-coupled interferometry (GCI), we identified six sybodies exhibiting affinities against the RBD ranging from 24 – 178 nM (Fig. 1A, Fig. S1A, Table S1). In this study, we focus on sybodies Sb#15 and Sb#68 (Fig. 1A), that can simultaneously bind the immobilized S-protein (Fig. 1B) and exhibit affinities of 12 nM and 9 nM, respectively, when probing against the entire spike protein stabilized by two prolines (S-2P) (Fig. 1A).

**Figure 1.**
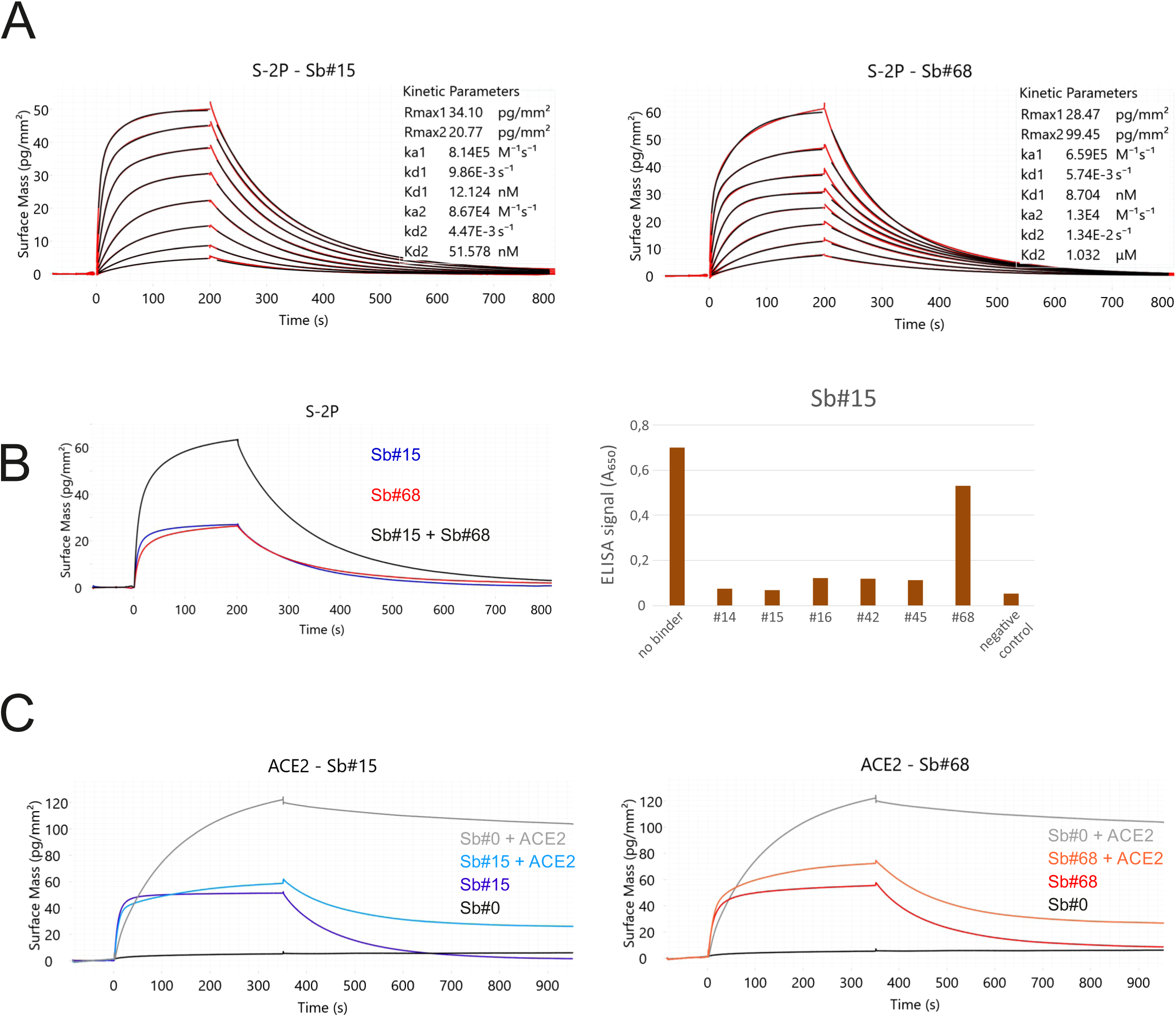
Sybodies Sb#15 and Sb#68 bind non-overlapping epitopes on the S-protein, and inhibit ACE2 binding. (**A**) Affinity determination of Sb#15 and Sb#68 against the immobilized S-protein (S-2P) using GCI. The data were fitted using a heterogeneous ligand model. (**B**) *left*, GCI epitope-binning experiment showing Sb#15 (blue), Sb#68 (red) and their combination (black) against immobilized S-protein (S-2P). Since both sybodies were present at saturating concentrations, the increased amplitude is indicative of simultaneous binding. Right, ELISA experiment confirming dual-binding of Sb#15 and Sb#68. Myc-tagged Sb#15 was immobilized on an anti-myc antibody-coated ELISA plate, followed by exposure of biotinylated RBD which was pre-mixed with tag-less sybodies (indicated on the x-axis). (C) Competition of sybodies and ACE2 for S-protein binding, investigated by GCI. Biotinylated S-protein was immobilized on the GCI sensor and then Sb#15 (200 nM, left), or Sb#68 (200 nM, right) were injected alone or premixed with human ACE2 (100 nM). Sb#0 represents a non-randomized control sybody.

To investigate whether Sb#15 and/or Sb#68 could block the interaction between the S-protein and ACE2, we performed an ACE2 competition experiment using GCI. To this end, S-protein was coated on a GCI chip and Sb#15 (200 nM), Sb#68 (200 nM) as well as a non-randomized convex sybody control (Sb#0, 200 nM) were injected alone or together with ACE2 (100 nM) to monitor binding (Fig. 1C). Indeed, Sb#0 did not bind when injected alone and consequently did not disturb ACE2 binding when co-injected. Conversely, both Sb#15 and Sb#68 were found to dominate over ACE2 in the association phase during co-injection, and the resulting curves are highly similar to what was observed when these two sybodies were injected alone. This experiment demonstrated that Sb#15 and Sb#68 compete with ACE2 for access to its binding site on the S-protein.

Having established that Sb#15 and Sb#68 could bind the S-protein and block ACE2 association in vitro, we next asked whether these sybodies (and the other four high affinity sybodies analyzed in Fig. S1) could inhibit the SARS-CoV-2 fusogenic machinery in viral neutralization assays. With the exception of Sb#42 and Sb#45, all sybodies neutralized vesicular stomatitis viruses (VSV) that were pseudotyped with SARS-CoV-2 S-protein [17], with IC_50_ values of 2.8 µg/ml (178 nM) and 2.3 µg/ml (138 nM), for Sb#15 and Sb#68, respectively (Fig. 2A, Table 1). Since Sb#15 and Sb#68 can bind simultaneously to full-length spike protein, we mixed Sb#15 and Sb#68 together to investigate potential additive or synergistic neutralizing activity of these two independent sybodies. Indeed, consistent with the binding assays, the simultaneous presence of both sybodies resulted in improved neutralization profiles with IC_50_ values reaching 1.7 µg/ml (53 nM). In addition to the individual sybodies, we also explored potential avidity effects of sybodies genetically fused to human IgG1 Fc domains. The respective homodimeric sybody-Fc constructs exhibited VSV pseudotype IC_50_ values of 1.2 µg/ml (15.5 nM) and 3.9 µg/ml (49.6 nM) for Sb#15 and Sb#68, respectively (Fig. 2B, Table 1). This improvement in VSV neutralization potency suggests that the bivalent arrangement of the Fc fusion constructs resulted in a discernible avidity effect. For neutralization of live SARS-CoV-2 we employed a plaque reduction assay and confirmed that both sybodies successfully inhibited cell entry by active SARS-CoV-2, with ND_50_ values of 37.4 µg/ml (2380 nM) for Sb#15 and 34.6 µg/ml (2070 nM) for Sb#68, but no neutralization by the other four sybodies (Table 1). The approximately 10-fold discrepancy in neutralization efficacies measured using either live SARS-CoV-2 virus or pseudotyped VSV may reflect slight differences in viral physiology (variation of incorporated spikes per viral particle) or could owe to the different assay methods (luciferase emission versus plaque reduction determination). Collectively, these data highlight the successful discovery of a pair of sybodies (Sb#15 and Sb#68) that bind simultaneously to the spike RBD, compete with ACE2 interaction, and neutralize viral infection in vitro.

**Figure 2.**
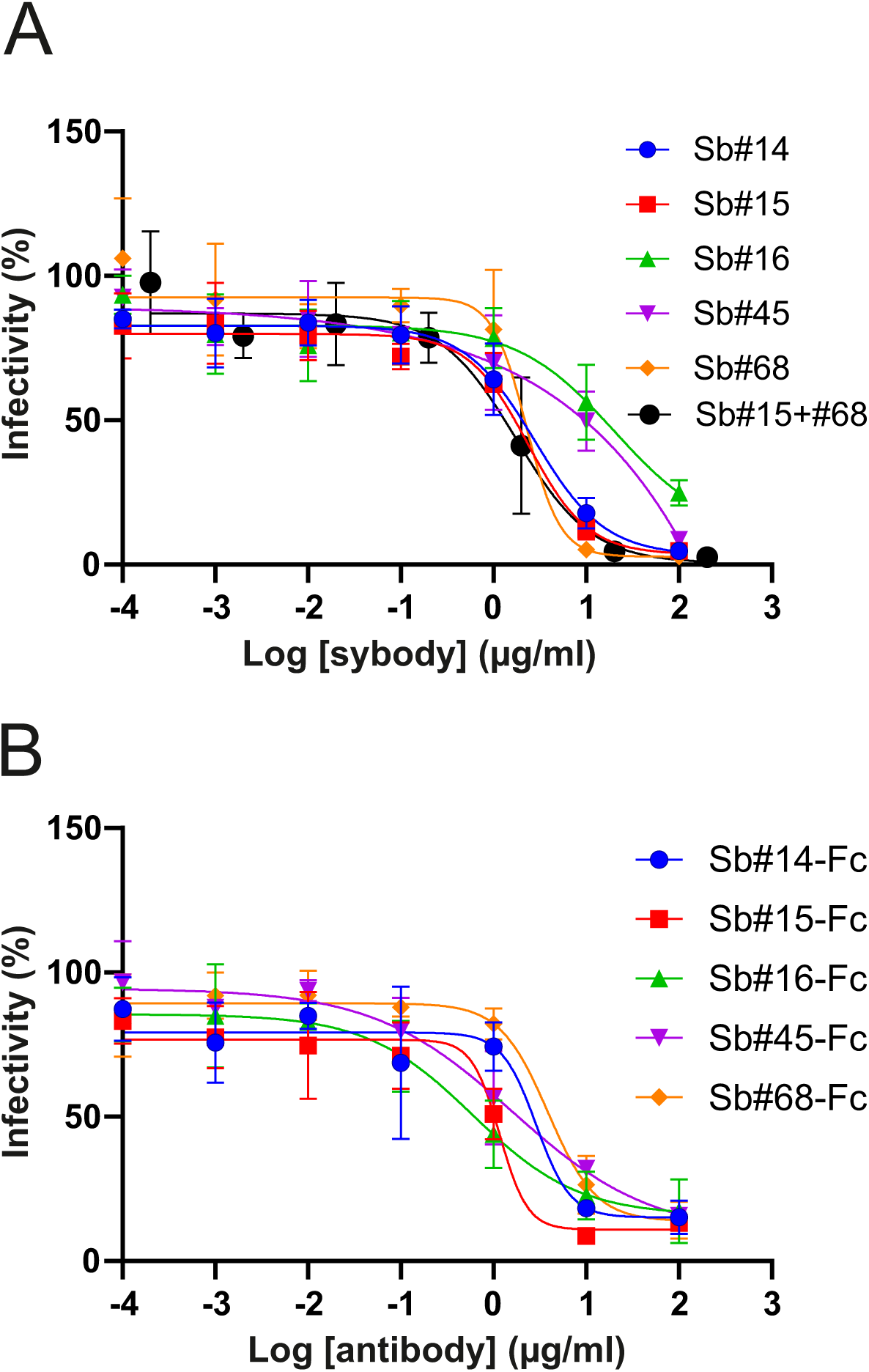
Neutralization of viral entry using pseudotyped VSVs. (**A**) Neutralization assay using VSVΔG pseudotyped with wildtype Spike-protein. Relative infectivity in response to increasing sybody concentrations was determined. The black curve shows data when a mixture of Sb#15 and Sb#68 was added. (**B**) Same assay as in (**A**) with sybodies fused to human Fc to generate bivalency. Error bars correspond to standard deviations of three biological replicates.

**Table 1.** summary of neutralization assay results

| Binders | SARS-CoV-2 pseudovirus |  | Live SARS-CoV-2 |  |
| --- | --- | --- | --- | --- |
|  | IC<sub>50</sub> (µg/ml) | IC<sub>50</sub> (nM) | ND<sub>50</sub> (µg/ml) | ND<sub>50</sub> (nM) |
| Sb#14 | 2.8 | 178.3 | nn | nn |
| Sb#15 | 2.3 | 146.5 | 37.4 | 2380 |
| Sb#16 | 20 | 1250 | nn | nn |
| Sb#68 | 2.3 | 137.7 | 34.6 | 2070 |
| Sb#15+Sb#68 | 1.7 | 52.5 | nd | nd |
| GS4 | 0.02 | 0.7 | 0.79 | 25.9 |
| Tripod-GS4r | 0.01 | 0.08 | nd | nd |
| Sb#14-Fc | 2.9 | 37.8 | nd | nd |
| Sb#15-Fc | 1.2 | 15.5 | nd | nd |
| Sb#16-Fc | 0.6 | 7.8 | nd | nd |
| Sb#45-Fc | 1.6 | 20.3 | nd | nd |
| Sb#68-Fc | 3.9 | 49.6 | nd | nd |

*nn: non-neutralizing*
*nd: not determined*

### Structural basis of Sb#15 and Sb#68 neutralizing activity

To gain structural insights into how Sb#15 and Sb#68 recognize the RBD and neutralize viruses, we performed single particle cryo-EM analysis of purified sybody-spike protein complexes. Three cryo-EM datasets were collected, allowing a glimpse of the spike protein either simultaneously bound to both sybodies, or associated to Sb#15 or Sb#68 alone (Fig. 3, Fig. S2-7, Table S2). The highest resolution was obtained for the spike protein in complex with both sybodies (Fig. 3, Fig. S2, Fig. S5), whereas structures with the individual sybodies were determined based on fewer particles and mainly served to unambiguously assign the binding epitopes of Sb#15 (Fig. S3, Fig. S6) and Sb#68 (Fig. S4, Fig. S7). Analysis of the spike/Sb#15/Sb#68 particles after 3D classification revealed that the spike protein adopts two distinct conformations (Fig. S2). The first conformation (30% of particles) has a three-fold symmetry, with three RBDs in the *up* conformation (*3up*) and two sybodies bound to each of the RBDs, confirming that Sb#15 and Sb#68 bind simultaneously (Fig. 3A, Fig. S2F and Fig. S5A).

**Figure 3.**
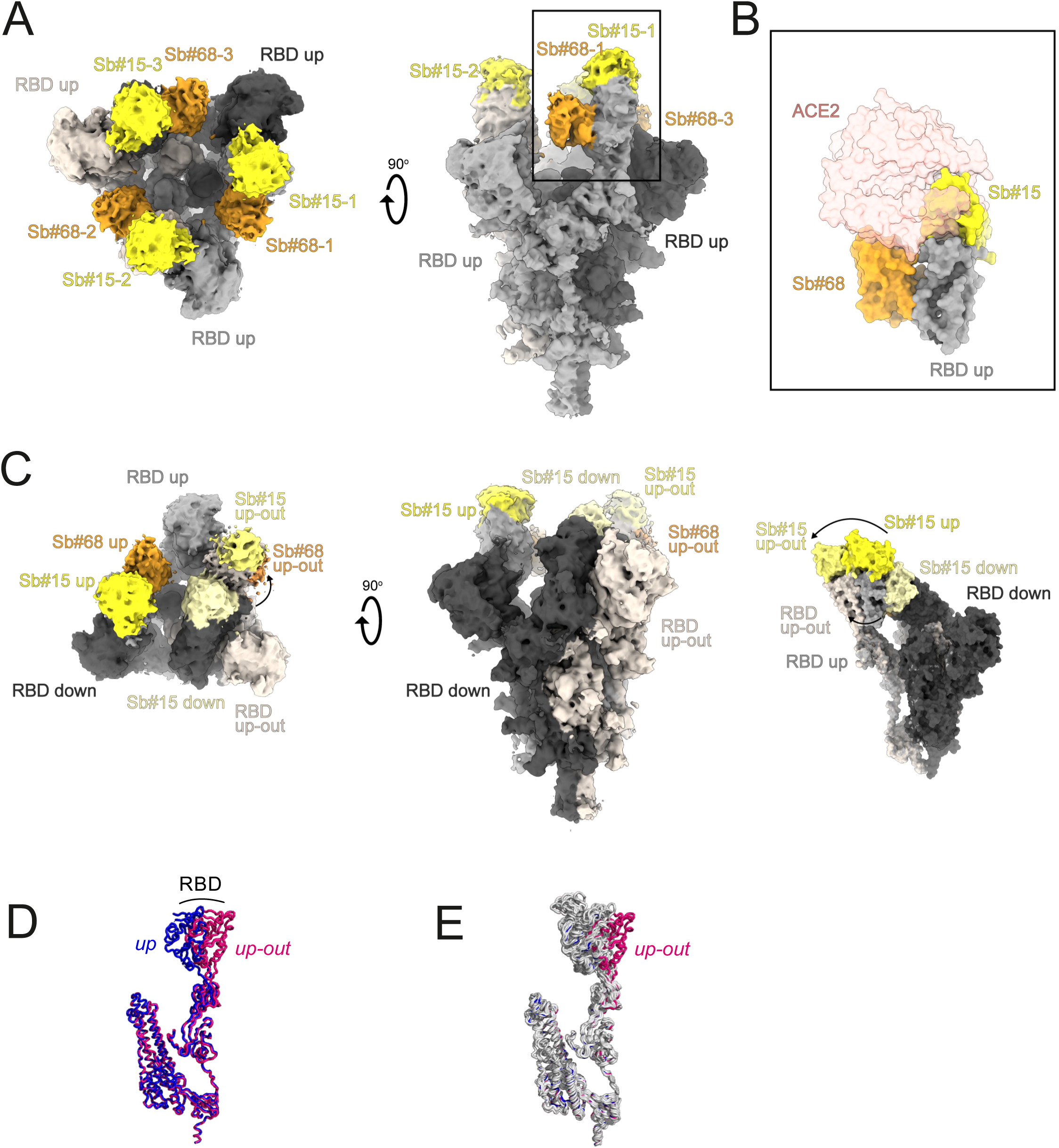
Cryo-EM maps of S-2P spike in complex with Sb#15 and Sb#68. (**A**) Cryo-EM map of S-2P with both Sb#15 and Sb#68 bound to each RBD adopting a symmetrical 3*up* conformation. (**B**) Close-up view showing that ACE2 binding to RBD (PDB ID: 6M0J) is blocked by bound Sb#15 and by a steric clash with Sb#68. (**C**) Cryo-EM map of S-2P with the three RBDs adopting an asymmetrical 1*up*/1*up-out*/1*down* conformation. Sb#15 is bound to all three RBDs, while Sb#68 is only bound to the *up* and *up-out* RBD. Final maps blurred to a B factor of -30 Å were used for better clarity of the less resolved RBDs and sybodies. Spike protein is shown in shades of grey, Sb#15 in yellow and Sb#68 in orange. (**D**) Alignment of structural models for the *up* (blue) and *up-out* (magenta) spike conformations. For clarity, only monomers are shown. (**E**) The *up-out* RBD conformation is unique among reported spike structures. Superposition of the aligned models (**D**) with 15 published structures of spike monomers showing up-RBDs (grey). PDB identifiers of aligned structures: 6VSB, 6VYB, 6XKL, 6ZGG, 6ZXN, 7A29, 7B18, 7CHH, 7CWT, 7DX9, 7JWB, 7LWW, 7M6F, 7N0H, 7N1V.

Although the global resolution of the spike protein in complex with both sybodies is around 3 Å, the local resolution of the RBDs with bound sybodies was only in the range of 6-7 Å, presumably due to conformational flexibility (Fig. S2). Therefore, we did not build a full model for Sb#15, but instead fitted a homology model into the density. In contrast, a crystal structure of Sb#68 in complex with the RBD had recently been determined at a resolution of 2.6 Å (PDB: 7KLW) [18], and therefore we used this high resolution structure of Sb#68 to fit it into the cryo-EM density.

Sb#15 binds to the top of the RBD. Its binding epitope roughly consists of two regions (residues 444-448 and 491-507) and thereby strongly overlaps with the ACE2 binding site (Fig. 3B). In contrast, Sb#68 binds to the side of the RBD (Fig. S4 and S7D-E) and recognizes a conserved “cryptic” epitope [8, 19] clearly distinct from the ACE2 interaction site, which includes residues 369-385 and R408 and is buried if the RBD is in its *down* conformation. Although the binding epitope of Sb#68 is clearly distinct from the one of ACE2, there would be a steric clash between the Sb#68 backside loops and ACE2, if ACE2 docks to the RBD (Fig. 3B). This accounts for Sb#68’s ability to compete with ACE2 as evident from GCI analyses (Fig. 1C).

The second resolved conformation (20 % of particles) of the spike/Sb#15/Sb#68 complex is asymmetric, with the RBDs in three distinct states, and was obtained at a global resolution of 3.3 Å (Fig. 3C, Fig. S2C, G and Fig. S5B). In this conformational state, each RBD was bound to Sb#15, whereas only two RBDs were associated with Sb#68-attributable cryo-EM density. One RBD was in the *up* conformation, having Sb#15 and Sb#68 bound in an analogous fashion as in the symmetric 3*up* structure. The orientation of this *up*-RBD closely superimposes with a variety of other reported cryo-EM structures (Fig. 3E). However, interestingly, a second RBD adopted a unique positioning that we term *up-out*. To our knowledge, this conformation has not been observed in prior S-protein structures, and therefore represents a novel RBD orientation (Fig. 3D and E). Notably, the density for Sb#68 was comparatively weak, indicating either high flexibility or a sub-stoichiometric occupancy. The novel *up-out* conformation appears to be caused by steric influences from the third RBD, which, in a *down* state and singularly bound to Sb#15, acts as a wedge that pushes the second RBD away from the three-fold symmetry axis and into its distinctive orientation (Fig. 3C).

Virtually the same asymmetric *1up/1up-out/1down* spike conformation was observed for the spike/Sb#15 complex, reinforcing our interpretation that wedging by Sb#15 is responsible for the outward movement of the second up-RBD (Fig. S6). However, according to our analysis, comprising only a limited number of images (Fig. S3D), Sb#15 alone was unable to induce the *3up* conformation, suggesting that adoption of the *3up* state requires the synergistic action of both sybodies to populate this symmetric conformation.

Finally, analysis of the spike/Sb#68 complex dataset revealed two distinct populations (Figure S4 and S7). The most abundant class showed an *1up/2dow*n conformation without sybody bound, which is identical to the one obtained for the spike protein alone [1, 20]. The second structure featured two RBDs in an *up* conformation with bound Sb#68. Density for the third RBD was very weak, presumably due to high intrinsic flexibility, hindering the interpretation of its exact position and conformation. We therefore refer to this conformation as an *2up/1flexible* state. Structural comparisons revealed that Sb#68 cannot access its epitope in the context of the *1up2dow*n conformation, due to steric clashes with the neighboring RBD (Fig. S7B). In order to bind, at least two RBDs need to be in the *up* conformation.

### Suppression of emergence of drug-resistant viruses by design of a biparatopic fusion construct

Biochemical and structural data provided evidence that fusing both sybodies may boost viral neutralization. To this end, Sb#15 and Sb#68 were genetically fused via a flexible (GGGGS)_4_ linker (Fig. 4A). The resulting purified bispecific Sb#15-(GGGGS)_4_-Sb#68 construct, designated as GS4, displayed a ≥40-fold increase in binding affinity for the S-protein (apparent Kd ≈ 0.3 nM), relative to either Sb#15 or Sb#68 alone (Fig. 4B). Strikingly, the neutralization potency of GS4 was increased by ≥100-fold over the individual binders, for both pseudotyped VSV (IC_50_ = 0.02 µg/ml; 0.7 nM) as well as for live SARS-CoV-2 (ND_50_ = 0.79 µg/ml; 25.9 nM) (Fig. 4C, Table 1), which suggest a highly synergistic inhibition mechanism mediated by the fusion molecule.

**Figure 4.**
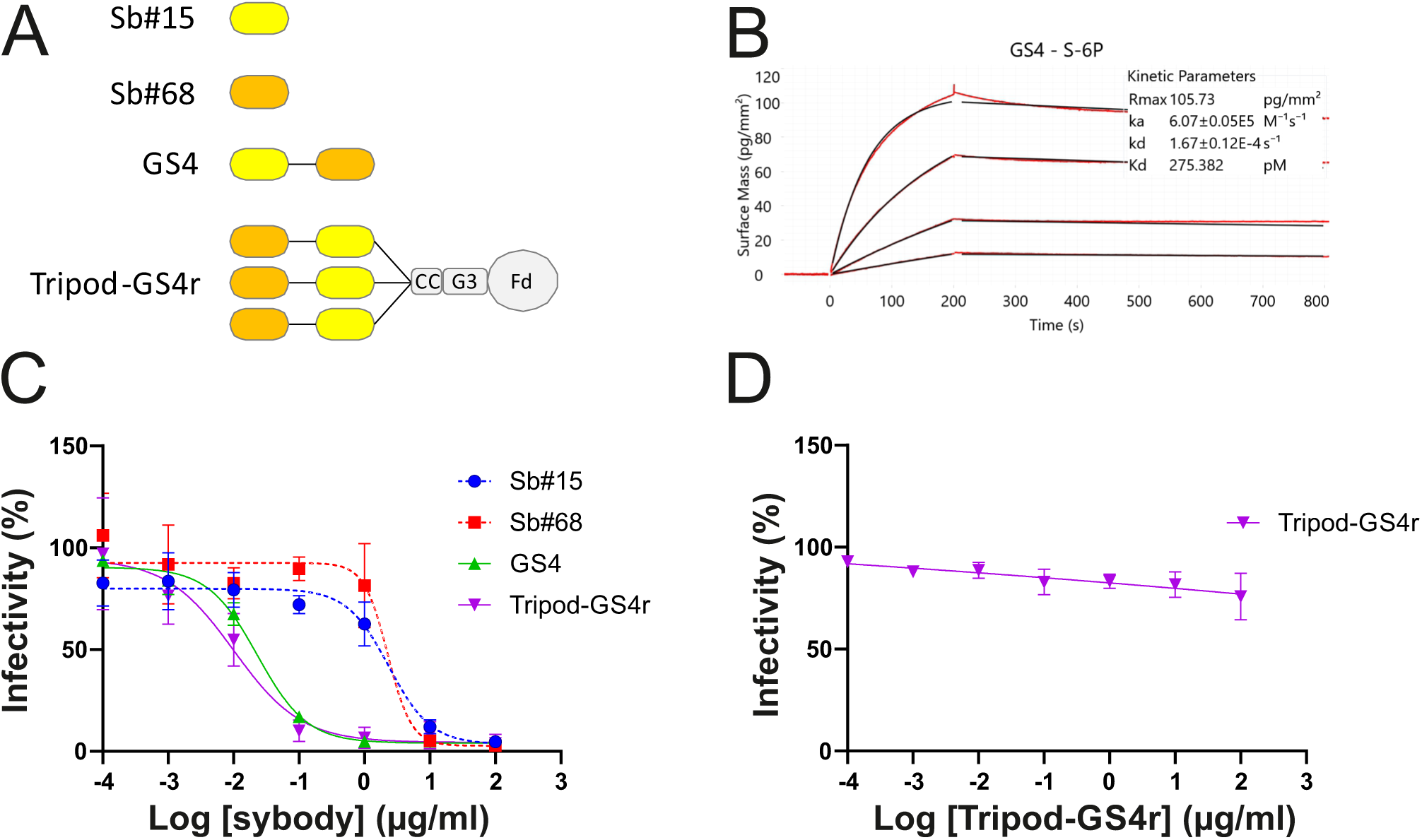
Fusion of Sb#15 and Sb#68 into biparatopic and multivalent molecules strongly enhance binding affinity and neutralization. (**A**) Schematic drawing of biparatopic GS4 and multivalent Tripod-GS4r constructs. (**B**) Affinity determination of GS4 against immobilized S-protein using GCI. (**C**) Neutralization assay using VSVΔG pseudotyped with wildtype Spike-protein. Relative infectivity was determined in response to increasing GS4 or Tripod-GS4r concentration. The corresponding neutralization data for isolated Sb#15 and Sb#68 are provided as reference. (**D**) Control experiment using VSVΔG pseudotyped with VSV-G. Infectivity was not affected byTripod-GS4r. Error bars correspond to standard deviations of three biological replicates.

We next asked whether simultaneous targeting of two spatially-distinct epitopes within the RBD would provide any advantages towards mitigating the development of escape mutants. By employing a replication-competent VSV^GFP^-SARS-CoV-2-S chimera [21] and a reported strategy to generate viral escape [22], we observed no resistant viruses in GS4-treated cells, whereas escape mutants emerged rapidly in the singular presence of either Sb#15 or Sb#68 (Fig. 5A, Table 2). Among the identified mutations from Sb#15- or Sb#68-treated cells, two were selected (Q493R for Sb#15 and P384H for Sb#68) and introduced into isolated RBDs (for binding kinetics measurements) and full-length S-protein (for neutralization determination). Indeed, our cryo-EM structures as well as a recently determined crystal structure of the RBD/Sb#68 complex were suggestive for a critical impact of both mutations on the respective epitope/paratope interfaces (Fig. 5A). Sb#15 and Sb#68 exhibited reduced binding with RBD-Q493R and RBD-P384H, respectively, although this attenuation was considerably more pronounced in case of the Sb#15/RBD-Q493R interaction (Fig. 5B). This correlated with the severely reduced neutralization efficacy by Sb#15 or Sb#68 against VSV expressing the corresponding adapted S-protein escape variants (Fig. 5C, Table 3). A previous study revealed that both mutations were neither significantly affecting ACE2 binding nor the overall expression profile, as revealed by a deep mutational scanning approach [23]. Finally, the bispecific fusion construct GS4 showed favorable binding kinetics and neutralization profiles in the presence of either individual mutation (Fig. 5B-C), supporting the hypothesis that simultaneously targeting multiple epitopes effectively mitigates evolutionary viral adaptation.

**Figure 5.**
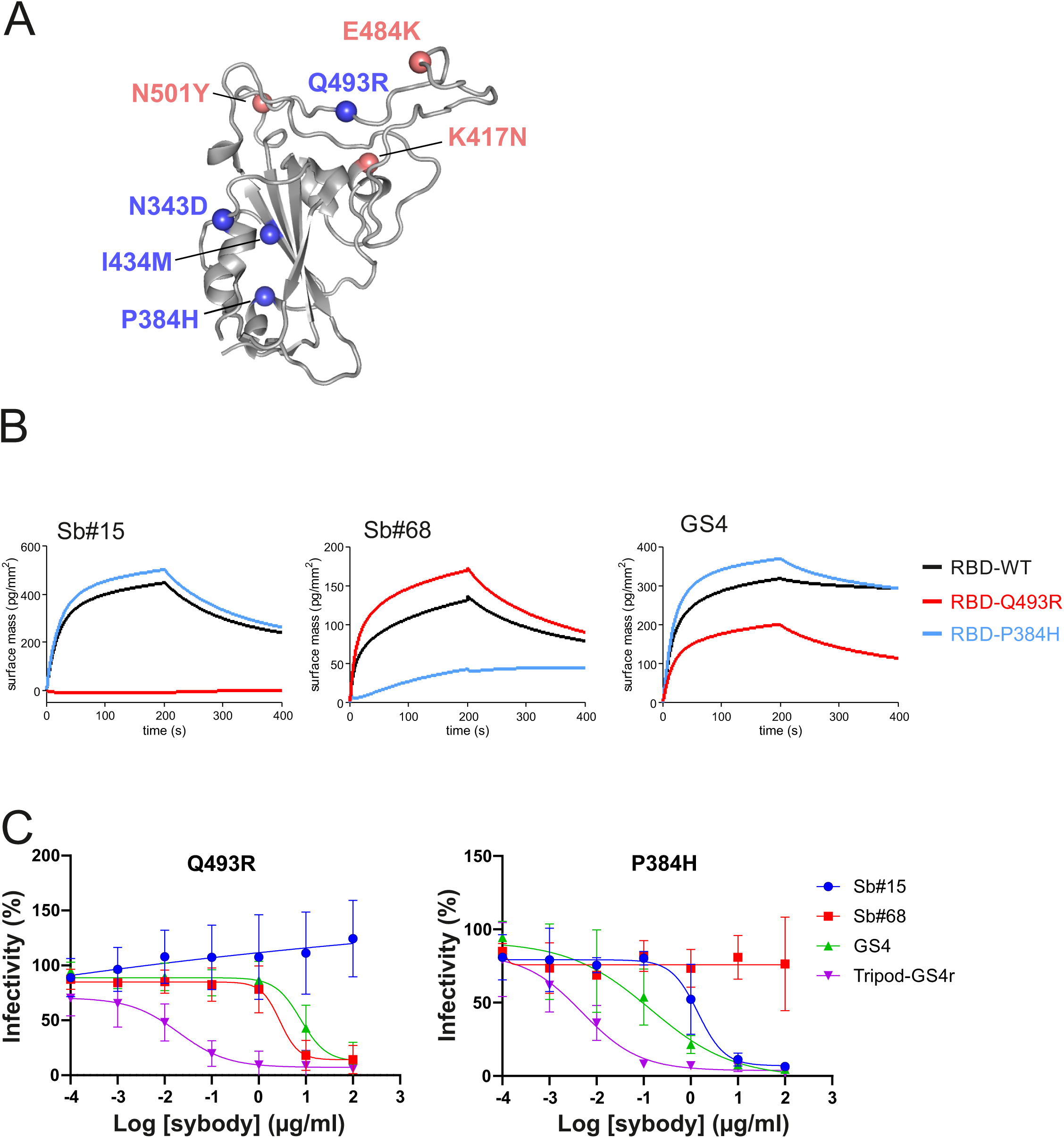
The bispecific construct GS4 mitigates emergence of novel escape mutants. (**A**) Structural context of RBD escape mutants resulting from adaptation experiments in the presence of either Sb#15 or Sb#68 alone (salmon spheres). Globally-circulating variants of concern are shown as blue spheres. (**B**) GCI-based kinetic analysis of the purified RBD bearing either no mutation (WT), or identified escape mutations Q493R or P384H. Left, middle, and right plots correspond to immobilized Sb#15, Sb#68, or GS4, respectively. (**C**) Neutralization assay using VSVΔG pseudotyped with Spike-protein containing the Q493R or P384H mutation, respectively. Relative infectivity in response to increasing binder concentrations was determined. Error bars correspond to standard deviations of three biological replicates.

**Table 2.**
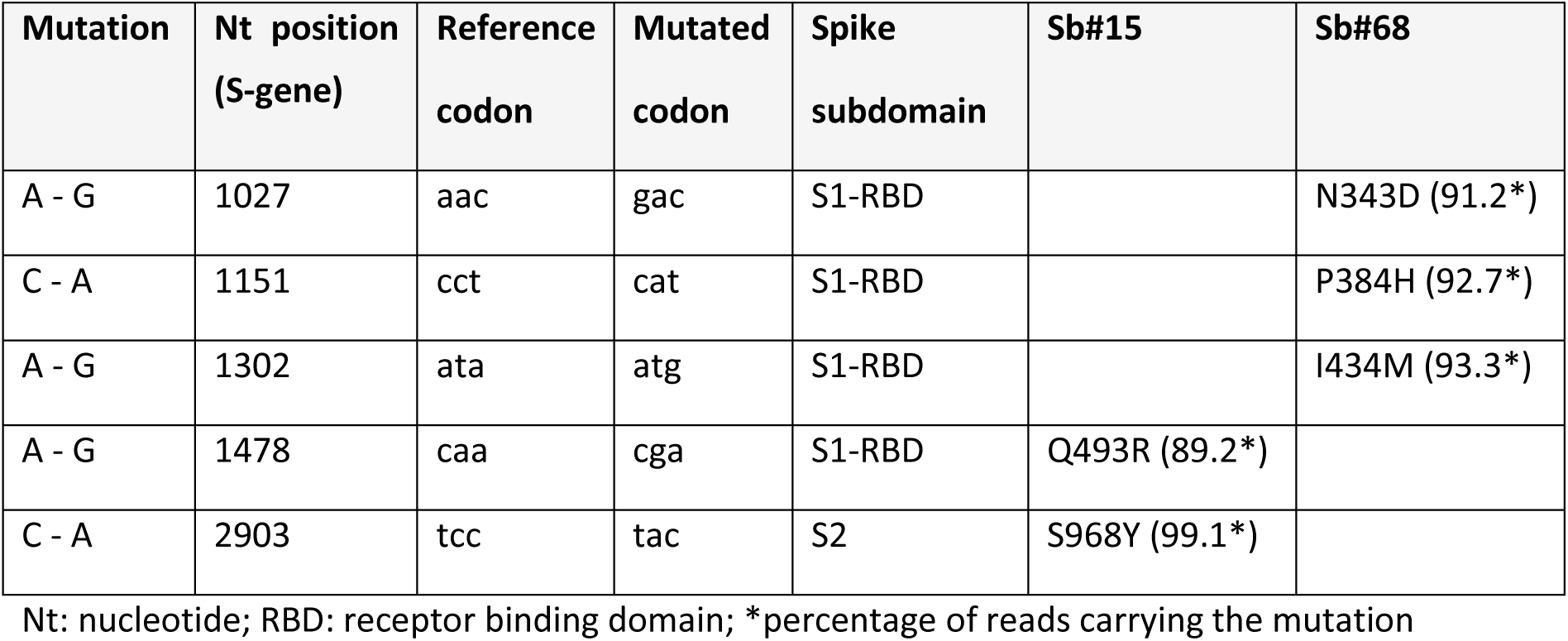
summary of missense escape mutations

| Mutation | Nt position (S-gene) | Reference codon | Mutated codon | Spike subdomain | Sb#15 | Sb#68 |
| --- | --- | --- | --- | --- | --- | --- |
| A - G | 1027 | aac | gac | S1-RBD |  | N343D (91.2*) |
| C - A | 1151 | cct | cat | S1-RBD |  | P384H (92.7*) |
| A - G | 1302 | ata | atg | S1-RBD |  | I434M (93.3*) |
| A - G | 1478 | caa | cga | S1-RBD | Q493R (89.2*) |  |
| C - A | 2903 | tcc | tac | S2 | S968Y (99.1*) |  |

691 Nt: nucleotide; RBD: receptor binding domain; \*percentage of reads carrying the mutation

**Table 3.** summary of neutralization assay results against specific spike mutants

|  | WT | B.1.1.7 (alpha) | RBD mutants |  |  |  | Escape mutants |  |
| --- | --- | --- | --- | --- | --- | --- | --- | --- |
|  |  |  | K417N | E484K | N501Y | KEN | Q493R | P384H |
| **15** | 147 | 1045 | >5000 | 841 | 1026 | >5000 | >5000 | 89 |
| **68** | 138 | 299 | 84 | 156 | 162 | 108 | 168 | >5000 |
| **GS4** | 0.7 | 16 | 7 | 1 | 7 | 436 | 256 | 5 |
| **Tripod-GS4r** | 0.08 | 0.09 | 0.04 | 0.11 | 0.12 | 0.08 | 0.16 | 0.04 |

IC<sub>50</sub> (nM)

### Activity of sybodies and multivalent fusion constructs against globally circulating SARS-CoV-2 variants of concern

We next investigated the efficacy of Sb#15, Sb#68 and the GS4 fusion construct against spike proteins harboring key mutations of the B.1.1.7 (alpha) and B.1.351 (beta) SARS-CoV-2 variants of concern. With regard to the RBD, S-alpha carries the N501Y substitution, whereas S-beta harbors the combination of K417N, E484K and N501Y mutations (Fig. 5A).

Consistent with the three mutations mapping at, or close to, the RBD/Sb#15 interface (K417N, E484K and N501Y), Sb#15 interacted with all single-mutation variants with reduced affinity, which was most pronounced for RBD-K417N (Fig. 6A). Compared to the individual mutations, the combined K417N/E484K/N501Y (KEN) triple mutant displayed a qualitatively additive effect with regards to reduced binding signal with Sb#15 (Fig. 6A). These binding kinetic phenotypes correlated with reduced neutralization efficacies for Sb#15 in pseudotyped VSV experiments with S-protein mutants (Fig. 6B, Table 3). In sharp contrast, Sb#68, which binds to the peripheral “cryptic epitope” of the RBD distant from the investigated KEN mutations, preserved proper binding kinetics and neutralization profiles against all spike variants (Fig. 6A, Table 3). Likely owing to this resilience of Sb#68 against the K417N/E484K/N501Y mutations, the binding affinities and neutralizing IC_50_ values of the bispecific GS4 molecule remained very favorable against these individual mutants (Fig 6B, Table 3). However, concerning the combined triple KEN mutant, although no significant impairment in the binding kinetics was recorded (Fig. 6A), the neutralizing IC_50_ values revealed that the bispecific construct GS4 was less potent than Sb#68 alone (about 400 nM versus 100 nM, respectively, Table 3). This suggests that, in the context of the KEN mutant, other potential factors that differ between purified spike protein and membrane-anchored spike protein in the context of VSVs influenced the overall neutralization profile of GS4.

**Figure 6.**
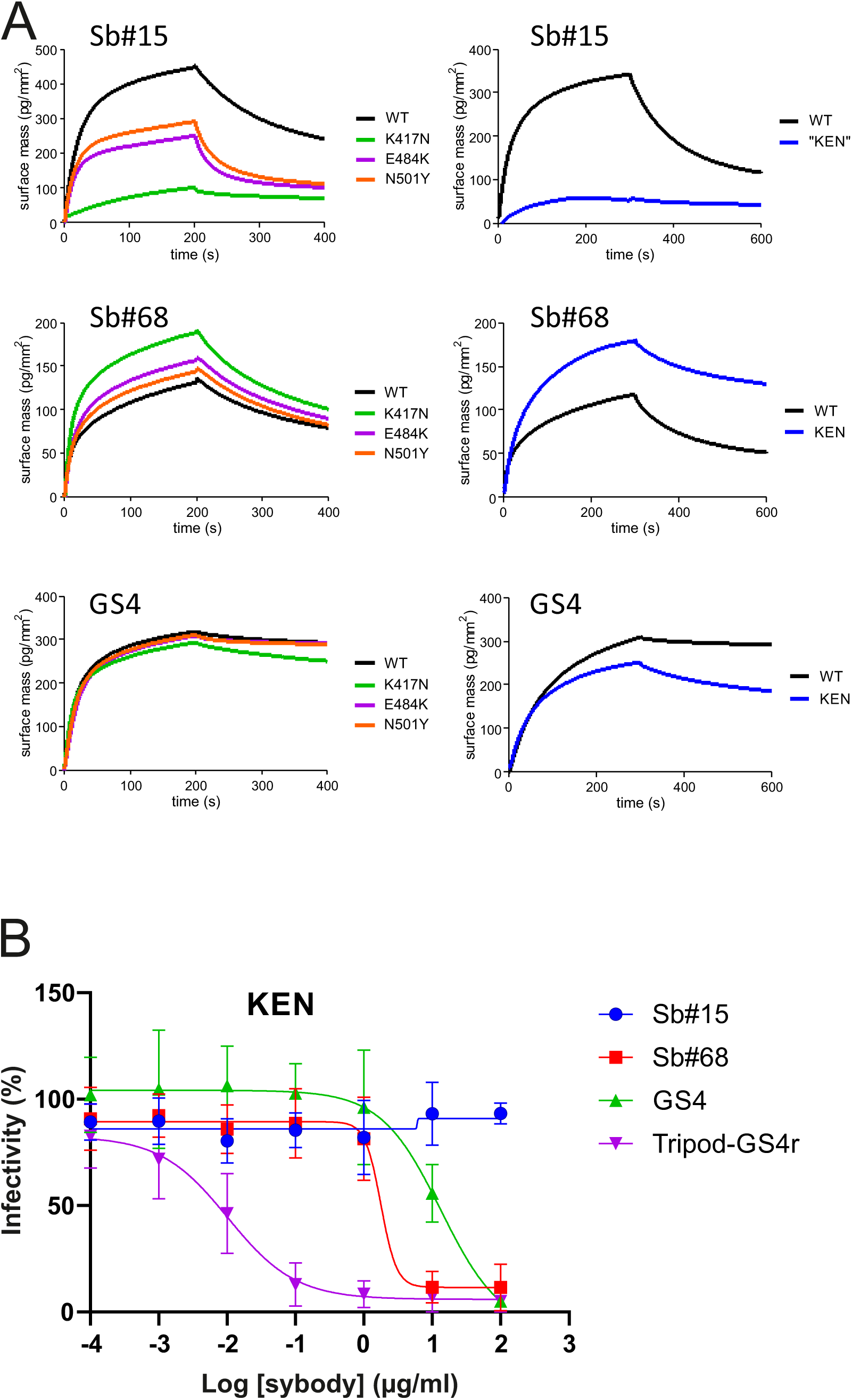
Affinity and neutralization by sybody constructs for variants of concern. (**A**) Semi-quantitative GCI analysis of interaction between RBDs carrying the individual K417N/E484K/N501Y mutations (left panels) or the combined triple KEN-RBD mutant (right panels). Sb#15, Sb#68 and GS4 were immobilized via a biotinylated Avi-tag and the RBD variants were injected. (**B**) Neutralization assay using VSVΔG pseudotyped with Spike-protein containing the triple KEN (beta) mutations. Relative infectivity in response to increasing binder concentrations was determined. Error bars correspond to standard deviations of three biological replicates.

We finally determined whether engineering additional layers of multivalency into the bispecific GS4 molecule would provide further advantages in neutralization profiles. To this aim, we grafted trimerization motifs (foldon [24], GCNt [25] and a covalently linked trimeric peptide) to the C-terminal region of Sb#15 via a flexible (GGGGS)_4_ linker (Fig. 4A, see Materials and Methods for more details). Furthermore, the N-terminal domain of Sb#15 was fused to the C-terminal region of Sb#68 via a second (GGGGS)_4_ linker. Of note, the position of both sybodies were reversed as compared to their orientation in GS4; this was based on the observed binding orientation of the sybodies in the context of our cryo-EM structures. This covalently-trimerized construct, termed Tripod-GS4r, was confirmed to undergo covalent multimerization that was reversible in reducing conditions (Fig. S8). Neutralization assays were conducted with VSV pseudotyped with wildtype spike or the spike protein harboring key mutations of the alpha or beta SARS-CoV-2 variants of concern (Fig. 4C, Fig. 6B). Remarkably, the recorded neutralizing IC_50_ values for tripod-GS4r were in the low picomolar range against viruses carrying all investigated spike mutations, including the triple KEN mutant (Table 3). Importantly, confirming specificity and lack of cytotoxicity, Tripod-GS4r did not neutralize control viruses pseudotyped with the non-cognate VSV-G spike glycoprotein (Fig. 4D).

## DISCUSSION

Since the start of the SARS-CoV-2 pandemic, multiple viral variants of concern have emerged, which threaten the progress made at the vaccination front [26]. Therefore, mitigation strategies that take into account the rapid evolution of the virus are urgently needed. Targeting multiple epitopes on the spike protein with large molecules can efficiently prevent viral escape, a strategy that was chosen by the pharmaceutical company Regeneron in bringing antibody cocktails on the market to treat COVID-19 patients [22, 27]. Nanobodies offer key advantages over conventional antibodies, in particular the ease of multimerization, inexpensive production and high protein stability. The latter simplifies logistics and facilitates development in an inhalable formulation [28, 29], thereby not only enabling direct delivery to nasal and lung tissues (two key sites of SARS-CoV-2 replication), but also offering the potential of self-administration.

Our study focused on a pair of sybodies, Sb#15 and Sb#68, which recognize two non-overlapping epitopes on the RBD. Both sybodies were found to compete with ACE2 binding. While the binding epitope of Sb#15 directly overlaps with the one of ACE2, this is not the case for Sb#68, which interferes with ACE2 through a steric clash at the sybody backside. Sb#15 and Sb#68 exhibited similar neutralization efficiencies, as well as a moderate synergistic effect in the virus neutralization test when both individual sybodies were mixed together. This synergy may be explained by the concerted action of the sybodies to compete with ACE2 docking via epitope blockage and steric clashing.

Cryo-EM analyses confirmed distinct binding epitopes for the two sybodies Sb#15 and Sb#68. Without sybodies, the spike protein predominantly assumes an equilibrium between the *3down* and the *1up2down* conformation [1, 20]. Upon addition of Sb#15, the conformational equilibrium was shifted towards an asymmetric *1up/1up-out/1down* state, whereas addition of Sb#68 favored an asymmetric state with RBDs adopting a *2up/1flexible* conformation. When added together, the sybodies synergized to stabilize two states: a predominant *3up* state, as well as the asymmetric *1up/1up-out/1down* state, thereby shifting the conformational equilibrium of the spike towards RBD conformations competent for ACE2 binding. These structural findings are reminiscent of a recent study, in which a pair of nanobodies isolated from immune libraries (VHH E and VHH V) was found to bind to similar epitopes as Sb#15 and Sb#68 [11]. However, in contrast to Sb#15 stabilizing the asymmetric *1up/1up-out/1down* state when added alone, the corresponding VHH E nanobody is exclusively bound to, and thereby stabilizes, the *3up* conformation. Hence, what is unique for our Sb#15/Sb#68 pair is its concerted action to shift the conformational equilibrium of the spike towards the *3up* state and its capability to trap the spike protein in an unusual *1up/1up-out/1down* conformation, which to the best of our knowledge has not been previously described.

Akin to the antibodies CR3022 and EY6A [19, 30] as well as a growing number of nanobodies [11, 31–33], the Sb#15/Sb#68 pair stabilized spike conformations with *2up* or *3up* RBDs. Thereby, the spike protein may be destabilized, resulting in the premature and unproductive transitions to the irreversible post-fusion state. This mechanism was dubbed “receptor mimicry” in a study on a neutralizing antibody S230, which only bound to up-RBDs and thereby triggered fusogenic conformational changes of SARS-CoV-1 spike [34]. In elegant experiments, König et al. could demonstrate that stabilization of the spike protein in its *3up* conformation by the addition of nanobodies VHH E and V indeed resulted in aberrant activation of the spike fusion machinery [11]. Hence, it is plausible to assume that our sybody pair inhibits SARS-CoV-2 infection and or/entry via such a receptor mimicry mechanism, in addition to blockage of ACE2 binding.

The binding epitope of Sb#68, also called the “cryptic” epitope [8], is highly conserved between SARS-CoV-1 and SARS-CoV-2. The same conserved epitope is also recognized by the human antibodies CR3022 (isolated from a SARS-CoV-1 infected patient and showing cross-specificity against SARS-CoV-2) and EY6A [19, 30] as well as the nanobody VHH-72, which had been originally selected against SARS-CoV-1 but was shown to cross-react with SARS-CoV-2 [35] (Fig. 7). Recent months have brought about a growing number of other nanobodies from immune and synthetic libraries whose epitopes overlap with Sb#68 [11, 31–33, 36, 37], suggesting that the cryptic epitope constitutes a preferred binding site for VHHs (Fig. 7C). The cryptic epitope is unchanged in currently circulating variants of concern, including the B.1.1.7 (alpha), B.1.351 (beta) and the B.1.617.2 (delta) lineages (Fig. 5A) and consequently, neutralization efficiency of Sb#68 is unaffected against these variants (Fig. 6B and C, Table 3).

**Figure 7.**
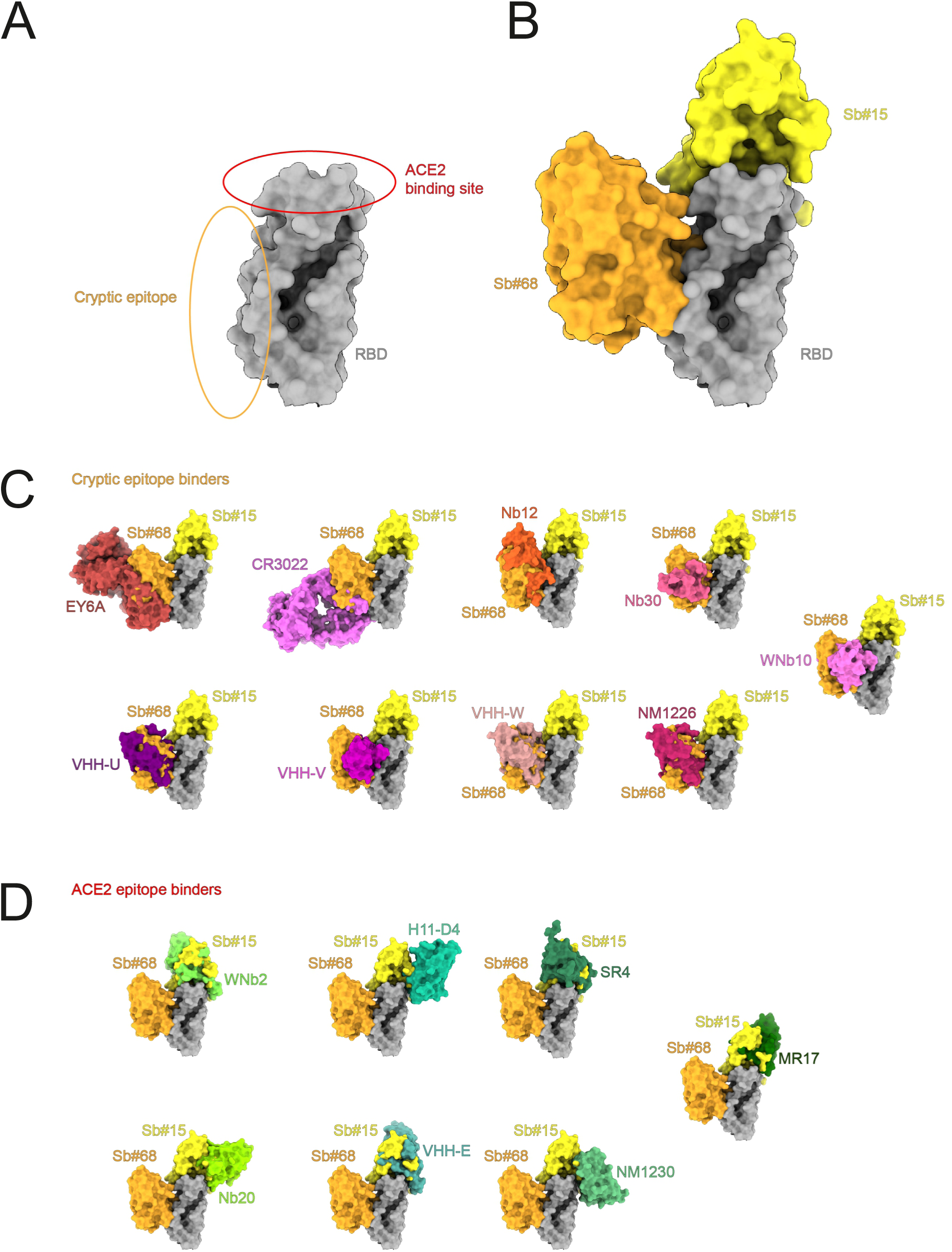
Antibody, nanobody and sybody binding to RBD epitopes. (**A**) Schematic representation of the ACE2 binding site and the cryptic epitope on the RBD. (**B**) Sb#15 binds to the top of RBD (the ACE2 epitope) and Sb#68 binds to the side of the RBD and recognizes a conserved cryptic epitope. (**C**) Superposition of the structures of binders recognizing the cryptic epitope: EY6A (PDB ID: 6ZCZ), CR3022 (PDB ID: 6W41), Nb12 (PDB ID:7MY3), Nb30 (PDB ID:7MY2), VHH-U (PDB ID: 7KN5), VHH-V (PDB ID: 7KN6), VHH-W (PDB ID: 7KN7), NM1226 (PDB ID: 7NKT), WNb10 (PDB ID: 7LX5). (**D**) Superposition of the structures of binders recognising the ACE2 epitope: WNb2 (PDB ID: 7LDJ), H11-D4 (PDB ID: 6YZ5), SR4 (PDB ID: 7C8V), Nb20 (PDB ID: 7JVB), VHH-E (7KN5), NM1230 (PDB ID: 7B27), MR17m(PDB ID: 7C8W). Structures are shown as surfaces.

Fusion of nanobodies via flexible linkers has emerged as a promising strategy to improve neutralization efficiencies by exploiting avidity effects. This potency-boosting procedure has been explored in the context of SARS-CoV-2 binders by either fusing up to three identical nanobodies (multivalency) [28, 31, 38], or by the structure-based design of biparatopic nanobodies [11, 39]. We exploited our structural data to first fuse Sb#15 and Sb#68 into the biparatopic GS4 construct, which resulted in a more than 100-fold gain of neutralization efficiency (Table 1). In a subsequent fusion step, the bi-paratopic GS4 construct was equipped with a trimerization domain (Tripod-GS4r), resulting in a further 10-fold boost in neutralization potential, thereby increasing the cumulative neutralization potency by a factor of around 1000 when compared to the single constituent nanobodies (Table 1). To our knowledge, engineering of such a “trimer-of-dimers” construct has not yet been attempted with anti-SARS-CoV-2 binder proteins. In addition to neutralizing virus entry via competition with receptor-binding and inducing premature activation of the fusion machinery, the unique multivalent structure of Tripod-GS4r may trigger clustering of neighboring spikes, thereby eventually deactivating multiple viral entry machineries simultaneously.

The ability of enveloped RNA viruses such as SARS-CoV-2 to rapidly develop resistance mutations is a crucial issue of consideration for the development of reliable therapeutics. Escape mutants indeed rapidly emerged when *in vitro* selection experiments were carried with single monoclonal antibodies or nanobodies targeting either the receptor-binding motif or the cryptic epitope alone [11, 22, 27]. It is interesting to note that the Q493R escape mutation we isolated for Sb#15 has been recently observed in a COVID-19 patient who received monoclonal antibody therapy [15]. Thus, an attractive strategy to potentially suppress mutational escape is to employ a cocktail of neutralizing antibodies binding to discrete epitopes of the spike. Accumulating evidence indeed demonstrate that rapid viral escape is not observed when experiments are performed either in the presence of a combination of neutralizing monoclonal antibodies/nanobodies or in the presence of biparatopic fusion constructs [11, 22, 27]. Furthermore, in addition to efficiently suppress mutational escape, our biparatopic molecules (*e.g.*, GS4 and Tripod-GS4r) consistently retained their neutralization capacity against pseudotyped VSV carrying spikes that harbored key mutations of common SARS-CoV-2 variants of concern. In particular, the trimerized biparatopic construct (Tripod-GS4r) exhibited ultra-potent neutralizing activity against all tested spike variants, with IC_50_ values of low picomolar range. Hence, our study provides evidence that combining multivalency and biparatopic nanobody fusion proteins represent a promising strategy to potentially generate therapeutic molecules with clinical relevance.

In conclusion, the rapid selection of sybodies [16] and their swift biophysical, structural and functional characterization, provide a foundation for the accelerated reaction to potential future pandemics. In contrast to a number of synthetic or naïve SARS-CoV-2 nanobodies from other libraries that required a post-selection maturation process to reach satisfactory affinities [28, 40–42], sybodies selected by us and by other labs [36, 43] exhibited affinities in the single and double digit nM range and where thus of similar affinity as nanobodies isolated from immune libraries using classical phage display [11, 37, 44]. Deep-mining of such sybody pools with our recently described flycode technology is expected to facilitate discovery of exceptional sybodies possessing very slow off-rates or recognizing rare epitopes [45]. Single domain antibodies and their derivative multi-component formats can be produced inexpensively and the biophysical properties of single domain antibodies make them feasible for development in an inhalable formulation, thereby offering the potential of self-administration. Hence, nanobodies show great promise to be used as prophylactic agents in the current or future pandemics.

## METHODS

### SARS-CoV-2 expression constructs and commercially-acquired proteins

For initial sybody selection experiments and binding affinity measurements, a gene encoding SARS-CoV-2 residues Pro330—Gly526 (RBD, GenBank accession QHD43416.1), downstream from a modified N-terminal human serum albumin secretion signal [46], was chemically synthesized (GeneUniversal). This gene was subcloned using FX technology [47] into a custom mammalian expression vector [48], appending a C-terminal 3C protease cleavage site, myc tag, Venus YFP[49], and streptavidin-binding peptide [50] onto the open reading frame (RBD-vYFP). A second purified RBD construct, consisting of SARS-CoV-2 residues Arg319—Phe541 fused to a murine IgG1 Fc domain was purchased from Sino Biological (RBD-Fc, Cat#: 40592-V05H). For kinetic interaction analysis of RBD escape mutants and sybodies, an RBD construct consisting of residues Arg319—Phe541, downstream from the native N-terminal SARS-CoV-2 secretion signal (Met1—Ser13) and appended with a C-terminal 10x-histidine tag (RBD-His), was cloned into a custom mammalian expression vector derived from pcDNA3.1 (ThermoFisher). Escape mutations P384H, K417N, E484K, Q493R, and N501Y were introduced into RBD-His using QuikChange site-directed mutagenesis. Expression plasmids harboring the prefusion ectodomain of the SARS-CoV2 spike protein (Met1—Gln1208), containing two or six stabilizing proline mutations (S-2P or S-6P, respectively) and a C-terminal foldon trimerization motif, HRV 3C protease cleavage site, and twin-strep tag, were a generous gift from Jason McLellan [1, 51]. Recombinant human ACE2 was purchased from mybiosource.com (Cat# MBS8248492). Recombinant Fc fusions of sybodies Sb#14, Sb#15, Sb#16, Sb#45, and Sb#68 were produced by Absolute Antibody.

### SARS-CoV-2 protein expression and purification

Suspension-adapted Expi293 cells (Thermo) were transiently transfected using Expifectamine according to the manufacturer protocol (Thermo), and expression was continued for 3–5 days in a humidified environment at 37°C, 8% CO_2_. Cells were pelleted (500*g*, 10 min), and culture supernatant was filtered (0.2 µm mesh size) before being incubated with the appropriate affinity chromatography matrix. For RBD-vYFP, NHS-agarose beads covalently coupled to the anti-GFP nanobody 3K1K[52] were used for affinity purification, and RBD-vYFP was eluted with 0.1 M glycine, pH 2.5, into tubes that were pre-filled with 1/10 vol 1M Tris (pH 9.0). Strep-Tactin®XT Superflow® (iba lifesciences) was used to pull down twin-strep-tagged S-2P or S-6P from culture supernatant, followed by elution with 50mM biotin. Ni-NTA beads were used for affinity purification of RBD-His, which was eluted with 300mM imidazole. All affinity-purified SARS-CoV-2 proteins were also subjected to size-exclusion chromatography using either a Superdex 200 Increase 10/300 GL column for RBD constructs, or a Superose 6 Increase 10/300 GL column for spike proteins.

### Sybody selections

Sybody selections, entailing one round of ribosome display followed by two rounds of phage display, were carried out as previously detailed with the three synthetic sybody libraries designated concave, loop and convex [16]. All targets were chemically biotinylated using NHS-Biotin (ThermoFisher #20217) according to the manufacturer protocol. Binders were selected against two different constructs of the SARS-CoV-2 RBD; an RBD-vYFP fusion and an RBD-Fc fusion. MBP was used as background control to determine the enrichment score by qPCR [16]. In order to avoid enrichment of binders against the fusion proteins (YFP and Fc), we switched the two targets after ribosome display. For the off-rate selections we did not use non-biotinylated target proteins as described[16] because we did not have the required amounts of purified target protein. Instead, we employed a pool competition approach. After the first round of phage display, all three libraries of selected sybodies, for both target-swap selection schemes, were subcloned into the pSb_init vector (giving approximately 10^8^ clones) and expressed in *E. coli* MC1061 cells. The resulting three expressed pools were subsequently combined, giving one sybody pool for each selection scheme. These two final pools were purified by Ni-NTA affinity chromatography, followed by buffer exchange of the main peak fractions using a desalting PD10 column in TBS pH 7.5 to remove imidazole. The pools were eluted with 3.2 ml of TBS pH 7.5. These two purified pools were used for the off-rate selection in the second round of phage display at concentrations of approximately 390 µM for selection variant 1 (competing for binding to RBP-Fc) and 450 µM for selection variant 2 (competing for binding to RBP-YFP). The volume used for off-rate selection was 500 µl, with 0.5% BSA and 0.05% Tween-20 added to pools immediately prior to the competition experiment. Off-rate selections were performed for 3 minutes. For identification of binder hits, ELISAs were performed as described [16]. 47 single clones were analyzed for each library of each selection scheme. Since the RBD-Fc construct was incompatible with our ELISA format due to the inclusion of Protein A to capture an α-myc antibody, ELISA was performed only for the RBD-vYFP (50 nM) and the M) and later on with the S-2P (25 nM). Of note, the three targets were analyzed in three separate ELISAs. As negative control to assess background binding of sybodies, we used biotinylated MBP (50 nM). 72 positive ELISA hits were sequenced (Microsynth, Switzerland).

### Expression and Purification of sybodies

The 63 unique sybodies were expressed and purified as described [16]. In short, all 63 sybodies were expressed overnight in *E.coli* MC1061 cells in 50 ml cultures. The next day the sybodies were extracted from the periplasm and purified by Ni-NTA affinity chromatography (batch binding) followed by size-exclusion chromatography using a Sepax SRT-10C SEC100 size-exclusion chromatography (SEC) column equilibrated in TBS, pH 7.5, containing 0.05% (v/v) Tween-20 (detergent was added for subsequent kinetic measurements). Six out of the 63 binders (Sb#4, Sb#7, Sb#18, Sb#34, Sb#47, Sb#61) were excluded from further analysis due to suboptimal behavior during SEC analysis (i.e. aggregation or excessive column matrix interaction).

### Generation of bispecific sybody fusions

To generate the bispecific sybodies (Sb#15-Sb#68 fusion with variable glycine/serine linkers), Sb#15 was amplified from pSb-init_Sb#15 (Addgene #153523) using the forward primer 5’-ATA TAT GCT CTT CAA GTC AGG TTC and the reverse primer 5’-TAT ATA GCT CTT CAA GAA CCG CCA CCG CCG CTA CCG CCA CCA CCT GCG CTC ACA GTC AC, encoding 2x a GGGGS motif, followed by a SapI cloning site. Sb#68 was amplified from pSb-init_Sb#68 (Addgene #153527) using forward primer 5’-ATA TAT GCT CTT CTT CTG GTG GTG GCG GTA GCG GCG GTG GCG GTA GTC AAG TCC AGC TGG TGG combined with the reverse primer 5’-TAT ATA GCT CTT CCT GCA GAA AC. The forward primers start with a SapI site (compatible overhang to Sb#15 reverse primer), followed by 2x the GGGGS motif. The PCR product of Sb#15 was cloned in frame with each of the three PCR products of Sb#68 into pSb-init using FX-cloning [47], thereby resulting in three fusion constructs with linkers containing 4x GGGGS motives as flexible linker between the sybodies (called GS4). The bispecific fusion construct GS4 was expressed and purified the same way as single sybodies [16].

### Construction, expression and purification of Tripod-GS4r

In order to engineer a trivalent GS4 molecule, we fused the following elements (from N- to C-terminus): Sb#68-(GGGGS)_4_-Sb#15-(GGGGS)_4_-CC-GCNt-Foldon-TST. CC is the trimeric CDV F protein (589-599) and contains two successive cysteine mutations (I595C and L596C) shown in the context of soluble measles virus F protein to stabilize the prefusion state [53]. GCNt is a trimerization motif that was previously described [25]. Foldon stems from fibritin [24]. TST denotes a C-terminal His/TwinStrepTag sequence for purification purposes. The Tripod-GS4r expression plasmid (3 mg) was sent to the Protein Production and Structure Core Facility of the EPFL (Switzerland) for expression (7 days in ExpiCHO cells). Subsequently, the protein was purified from 1 L of supernatant using a 5 mL StrepTtrapXT column (Cytavia) and eluted with 500 mM biotin (Cytivia).

### Dual-sybody competition ELISA

Purified sybodies carrying a C-terminal myc-His Tag (Sb_init expression vector) were diluted to 25 nM in 100 µl PBS pH 7.4 and directly coated on Nunc MaxiSorp 96-well plates (ThermoFisher #44-2404-21) at 4°C overnight. The plates were washed once with 250 µl TBS pH 7.5 per well followed by blocking with 250 µl TBS pH 7.5 containing 0.5% (w/v) BSA per well. In parallel, chemically biotinylated prefusion Spike protein (S-2P) at a concentration of 10 nM was incubated with 500 nM sybodies for 1 h at room temperature in TBS-BSA-T. The plates were washed three times with 250 µl TBS-T per well. Then, 100 µl of the S-2P-sybody mixtures were added to the corresponding wells and incubated for 3 min, followed by washing three times with 250 µl TBS-T per well. 100 µl Streptavidin-peroxidase polymer (Merck, Cat#S2438) diluted 1:5000 in TBS-BSA-T was added to each well and incubated for 10 min, followed by washing three times with 250 µl TBS-T per well. Finally, to detect S-2P bound to the immobilized sybodies, 100 µl ELISA developing buffer (prepared as described previously [16]) was added to each well, incubated for 1 h (due to low signal) and absorbance was measured at 650 nm. As a negative control, TBS-BSA-T devoid of protein was added to the corresponding wells instead of a S-2P-sybody mixture.

### Grating-coupled interferometry (GCI)

Kinetic characterization of sybodies binding onto SARS-CoV-2 spike proteins was performed using GCI on the WAVEsystem (Creoptix AG, Switzerland), a label-free biosensor. For the off-rate screening, biotinylated RBD-vYFP and ECD were captured onto a Streptavidin PCP-STA WAVEchip (polycarboxylate quasi-planar surface; Creoptix AG) to a density of 1300-1800 pg/mm^2^. Sybodies were first analyzed by an off-rate screen performed at a concentration of 200 nM (data not shown) to identify binders with sufficiently high affinities. The six sybodies Sb#14, Sb#15, Sb#16, Sb#42, Sb#45, and Sb#68 were then injected at increasing concentrations ranging from 1.37 nM to 1 μM (three-fold serial dilution, 7 concentrations) in 20 mM Tris pH7.5, 150 mM NaCl supplemented with 0.05 % Tween-20 (TBS-T buffer). Sybodies were injected for 120 s at a flow rate of 30 μl/min per channel and dissociation was set to 600 s to allow the return to baseline.

In order to determine the binding kinetics of Sb#15 and Sb#68 against intact spike proteins, the ligands RBD-vYFP, S-2P and S-6P were captured onto a PCP-STA WAVEchip (Creoptix AG) to a density of 750 pg/mm^2^, 1100 pg/mm^2^ and 850 pg/mm^2^ respectively. Sb#15 and Sb#68 were injected at concentrations ranging from 1.95 nM to 250 nM or 3.9 nM to 500 nM, respectively (2-fold serial dilution, 8 concentrations) in TBS-T buffer. Sybodies were injected for 200 s at a flow rate of 80 μl/min and dissociation was set to 600 s. In order to investigate if Sb#15 and Sb#68 bind simultaneously to the RBD, S-2P and S-6P, both binders were either injected alone at a concentration of 200 nM or mixed together at the same individual concentrations at a flow rate of 80 μl/min for 200 s in TBS-T buffer.

To measure binding kinetics of the three bispecific fusion construct GS4, S-6P was captured as described above to a density of 1860 pg/mm^2^ and increasing concentrations of the bispecific fusion constructs ranging from 1 nM to 27 nM (3-fold serial dilution, 4 concentrations) in TBS-T buffer at a flow rate of 80 μl/min. Because of the slow off-rates, we performed a regeneration protocol by injecting 10 mM glycine pH 2 for 30 s after every binder injection.

For ACE2 competition experiments, S-2P was captured as described above. Then Sb#15, Sb#68 or Sb#0 (non-randomized convex sybody control) were either injected individually or premixed with ACE2 in TBS-T buffer. Sybody concentrations were at 200 nM and ACE2 concentration was at 100 nM.

All sensorgrams were recorded at 25 °C and the data analyzed on the WAVEcontrol (Creoptix AG). Data were double-referenced by subtracting the signals from blank injections and from the reference channel. A Langmuir 1:1 model was used for data fitting with the exception of the Sb#15 and Sb#68 binding kinetics for the S-2P and the S-6P spike, which were fitted with a heterogeneous ligand model as mentioned in the main text.

### SARS-CoV-2 pseudovirus neutralization

Pseudovirus neutralization assays have been previously described [17, 35, 54]. Briefly, propagation-defective, spike protein-pseudotyped vesicular stomatitis virus (VSV) was produced by transfecting HEK-239T cells with SARS-CoV-2 Sdel 18 (SARS-2 S carrying an 18 aa cytoplasmic tail truncation) as described previously [55]. The cells were further inoculated with glycoprotein G *trans*-complemented VSV vector (VSV*ΔG(Luc)) encoding enhanced green fluorescence protein (eGFP) and firefly luciferase reporter genes but lacking the glycoprotein G gene [56]. After 1 h incubation at 37 °C, the inoculum was removed and the cells were washed once with medium and subsequently incubated for 24 h in medium containing 1:3000 of an anti-VSV-G mAb I1 (ATCC, CRL-2700^TM^). Pseudotyped particles were then harvested and cleared by centrifugation.

For the SARS-CoV-2 pseudotype neutralization experiments, pseudovirus was incubated for 30 min at 37 °C with different dilutions of purified sybodies, sybdody fusions or sybody-Fc fusions. Subsequently, S protein-pseudotyped VSV*ΔG(Luc) was added to Vero E6 cells grown in 96-well plates (25’000 cells/well). At 24 h post infection, luminescence (firefly luciferase activity) was measured using the ONE-Glo Luciferase Assay System (Promega) and Cytation 5 cell imaging multi-mode reader (BioTek).

### SARS-CoV-2 neutralization test

The serial dilutions of control sera and samples were prepared in quadruplicates in 96-well cell culture plates using DMEM cell culture medium (50 µL/well). To each well, 50 µL of DMEM containing 100 tissue culture infectious dose 50% (TCID_50_) of SARS-CoV-2 (SARS-CoV-2/München-1.1/2020/929) were added and incubated for 60 min at 37°C. Subsequently, 100 µL of Vero E6 cell suspension (100,000 cells/mL in DMEM with 10% FBS) were added to each well and incubated for 72 h at 37 °C. The cells were fixed for 1 h at room temperature with 4% buffered formalin solution containing 1% crystal violet (Merck, Darmstadt, Germany). Finally, the microtiter plates were rinsed with deionized water and immune serum-mediated protection from cytopathic effect was visually assessed. Neutralization doses 50% (ND_50_) values were calculated according to the Spearman and Kärber method.

### Generation of chimeric VSV\***Δ**G-S**_Δ_**_21_

Recently, we generated a recombinant chimeric VSV, VSV*ΔG(MERS-S), in which the VSV glycoprotein (G) gene was replaced by the full-length MERS-CoV spike protein [21]. VSV*ΔG(MERS-S) also encoded a GFP reporter which was expressed from an additional transcription unit located between the MERS-CoV spike and VSV L genes. In order to generate a chimeric VSV expressing the SARS-CoV-2 spike protein, the MERS-S gene in the antigenomic plasmid pVSV*ΔG(MERS-S) was replaced by a modified SARS-CoV-2 spike gene (Genscript, Piscattaway, USA) taking advantage of the flanking *Mlu*I and *BstE*II endonuclease restriction sites. The modified SARS-CoV-2 spike gene was based on the Wuhan-Hu-1 strain (GenBank accession number NC_045512) but lacked the region encoding the C-terminal 21 amino acids in order to enhance cell surface transport of the spike protein and its incorporation into the VSV envelope [57]. In addition, the modified spike gene contained the mutations R685G, H655Y, D253N, W64R, G261R, A372T, which have been previously reported to accumulate during passaging chimeric VSV-SARS-CoV-2-S on Vero E6 cells [57]. The amino acid substitution R685G is located in the S1/S2 proteolytic cleavage site and has been shown to reduce syncytia formation and to enhance virus titers [57].

The chimeric virus was rescued following transfection of cDNA according to a previously described protocol [58]. Briefly, BHK-21 cells were infected with a modified virus Ankara (MVA) expressing T7 bacteriophage RNA polymerase [59] using a multiplicity of infection of 1 focus-forming unit (ffu)/cell. Subsequently, the cells were transfected with the VSV antigenomic plasmid along with plasmids driving the T7 RNA polymerase-mediated transcription of the VSV N, P, and L genes. After 24 hours of incubation, the cells were trypsinized and seeded along with an equal number of Vero E6 cells and incubated for an additional 48 hours at 37°C. The expression of the GFP reporter in the cells was monitored by fluorescence microscopy. The recombinant virus was rescued from the supernatant of GFP-positive cells and passaged subsequently on Vero E6 cells. Following the fourth passage, virus was harvested from the cell culture supernatant and stored in aliquots at -70°C in the presence of 10% fetal bovine serum (FBS).

Infectious virus titers were determined on confluent Vero E6 cells grown in 96-well microtiter plates. The cells were inoculated in duplicate with 40 μl per well of serial 10-fold virus dilutions for 1 hour at 37°C. Thereafter, 160 μl of EMEM containing 1% methyl-cellulose was added to each well, and the cells were incubated for 24 hours at 37°C. The number of infectious foci was determined under the fluorescence microscope taking advantage of the GFP reporter and infectious virus titers were calculated and expressed as ffu/ml.

### Selection of VSV\***Δ**G-S**_Δ_**_21_ escape mutants

The selection of VSV*ΔG-S_Δ21_ mutants which escaped sybody-mediated inhibition was performed according to a recently described procedure [22]. Briefly, a total of 10^4^ ffu of the parental VSV*ΔG-S_Δ21_ were incubated with serially diluted sybodies prior to infection of Vero E6 cells that were grown in 24-well cell culture plates. Two days post infection, the cell culture supernatant from wells containing the highest antibody concentration which did not completely inhibit virus replication as monitored by GFP fluorescence was collected and subjected to a second round of selection on Vero E6 cells grown in 96-well microtiter plates in the presence of increasing sybody concentrations. Virus recovered after a third round of selection was used to infect Vero E6 cells grown in 6-well plates.

### NGS analysis of escape mutants

Vero E6 cells were lysed 24 hours post-infection with TRIZol reagent (Ambion, Life Technologies, Zug, Switzerland) and total RNA was subsequently isolated using a Direct-zol^TM^ RNA MicroPrep Kit (Zymo Research #R2060) according to the manufacturer’s protocol. The recommended DNase treatment was included. The quantity and quality of the extracted RNA was assessed using a Thermo Fisher Scientific Qubit 4.0 fluorometer with the Qubit RNA BR Assay Kit (Thermo Fisher Scientific, Q10211) and an Advanced Analytical Fragment Analyzer System using a Fragment Analyzer RNA Kit (Agilent, DNF-471), respectively. Prior to cDNA library generation, probe-based depletion of ribosomal RNA was performed on 500 ng of total RNA using a RiboCop rRNA Depletion Kit -Human/Mouse/Rat plus Globin (Lexogen #145.96) according to the producer’s protocol. Thereafter, the remaining RNA was used as input for a CORALL Total RNA-Seq Library Prep Kit (Lexogen #117.96) in combination with Lexogen workflow A unique dual indexes Set A1 (lexogen UDI12A_0001-0096) following the corresponding user guide (Lexogen document 117UG228V0200). The quantity and quality of the generated NGS libraries were evaluated using a Thermo Fisher Scientific Qubit 4.0 fluorometer with the Qubit dsDNA HS Assay Kit (Thermo Fisher Scientific, Q32854) and an Advanced Analytical Fragment Analyzer System using a Fragment Analyzer NGS Fragment Kit (Agilent, DNF-473), respectively. Pooled cDNA libraries were paired-end sequenced using a NovaSeq 6000 SP reagent kit v1.5, 300 cycles (illumina 20028402) on an Illumina NovaSeq 6000 instrument. The quality of the sequencing runs was assessed using illumina Sequencing Analysis Viewer (illumina version 2.4.7) and all base call files were demultiplexed and converted into FASTQ files using illumina bcl2fastq conversion software v2.20. The RNA quality-control assessments, generation of libraries and sequencing runs were performed at the Next Generation Sequencing Platform, University of Bern, Switzerland.

### Bioinformatic analysis

Reads were trimmed using trimmomatic [60] version 0.36, while setting the ILLUMINACLIP parameter to 2:30:10, SLIDINGWINDOW to 4:5 and MINLEN to 50. The parameter HEADCROP was set to 12 to remove the UMI sequence at the forward reads and low-quality bases due to random priming of the reverse read. Trimmed reads were aligned against the spike construct sequence (encoding for the mutations R685G, H655Y, D253N, W64R, G261R, A372T) by using minimap2 [61] with the short read mode (-x sr). Read duplicates were marked in the alignment file with samtools markdup [62]. Variants were called on each sample separately using gatk4 HaplotypCaller [63] with setting the option --sample-ploidy to 1 and --emit-ref-confidence to GVCF. The resulting GVCF files were combined into a single VCF file with gatk GenomicsDBImport and GenotypeGVCFs. The alleles and counts per allele were reported for each variant for each sample by using bcftools [62] query.

### Cryo-EM sample preparation and data collection

Freshly purified S-2P was incubated with a 1.3-fold molar excess of Sb#15 alone or with Sb#15 and Sb#68 and subjected to size exclusion chromatography to remove excess sybody. In analogous way, the sample of S-6P with Sb#68 was prepared. The protein complexes were concentrated to 0.7-1 mg ml^-1^ using an Amicon Ultra-0.5 mL concentrating device (Merck) with a 100 kDa filter cut-off. 2.8 μl of the sample was applied onto the holey-carbon cryo-EM grids (Au R1.2/1.3, 300 mesh, Quantifoil), which were prior glow discharged at 5 - 15 mA for 30 s, blotted for 1–2 s and plunge frozen into a liquid ethane/propane mixture with a Vitrobot Mark IV (Thermo Fisher) at 15 °C and 100% humidity. Samples were stored in liquid nitrogen until further use. Screening of the grid for areas with best ice properties was done with the help of a home-written script to calculate the ice thickness (manuscript in preparation). Cryo-EM data in selected grid regions were collected in-house on a 200-keV Talos Arctica microscope (Thermo Fisher Scientifics) with a post-column energy filter (Gatan) in zero-loss mode, with a 20-eV slit and a 100 μm objective aperture. Images were acquired in an automatic manner with SerialEM on a K2 summit detector (Gatan) in counting mode at ×49,407 magnification (1.012 Å pixel size) and a defocus range from −0.9 to −1.9 μm. During an exposure time of 9 s, 60 frames were recorded with a total exposure of about 53 electrons/Å^2^. On-the-fly data quality was monitored using FOCUS [64].

### Image processing

For the S-2P/Sb#15/ Sb#68 complex dataset, in total 14,883 micrographs were recorded. Beam-induced motion was corrected with MotionCor2_1.2.1 [65] and the CTF parameters estimated with ctffind4.1.13 [66]. Recorded micrographs were manually checked in FOCUS (1.1.0), and micrographs, which were out of defocus range (<0.4 and >2 μm), contaminated with ice or aggregates, and with a low-resolution estimation of the CTF fit (>5 Å), were discarded. 637,105 particles were picked from the remaining 12,454 micrographs by crYOLO 1.7.5 [39], and imported in cryoSPARC v2.15.0 [67] for 2D classification with a box size of 300 pixels. After 2D classification, 264,082 particles were imported into RELION-3.0.8 [68] and subjected to a 3D classification without imposed symmetry, where an ab-initio generated map from cryoSPARC low-pass filtered to 50 Å was used as reference. Two classes resembling spike protein, revealed two distinct conformations. One class shows a symmetrical state with all RBDs in an *up* conformation (3*up*) and both sybodies bound to each RBD (78,933 particles, 30%). In the asymmetrical class (52,839 particles, 20%) the RBDs adopt one *up*, one *up-out* and one *down* conformation (*1up/1up-out/1down)*, where both sybodies are bound to RBDs *up* and *up-out* state, while only Sb#15 is bound to the *down* RBD. The *3up* class was further refined with C3 symmetry imposed. The final refinement, where a mask was included in the last iteration, provided a map at 7.6 Å resolution. Six rounds of per-particle CTF refinement with beamtilt estimation and re-extraction of particles with a box size of 400 pixels improved resolution further to 3.2 Å. The particles were then imported into cryoSPARC, where non-uniform refinement improved the resolution to 3 Å. The asymmetrical *1up/1up-out/1down* was refined in an analogous manner with no symmetry imposed, resulting in a map at 7.8 Å resolution. Six rounds of per-particle CTF refinement with beamtilt estimation improved resolution to 3.7 Å. A final round of non-uniform refinement in cryoSPARC yielded a map at 3.3 Å resolution. Local resolution estimations were determined in cryoSPARC. All resolutions were estimated using the 0.143 cut-off criterion [69] with gold-standard Fourier shell correlation (FSC) between two independently refined half-maps [70]. The directional resolution anisotropy of density maps was quantitatively evaluated using the 3DFSC web interface (https://3dfsc.salk.edu) [71].

A similar approach was performed for the image processing of the S-2P/Sb#15 complex. In short, 2,235 micrographs were recorded, and 1,582 used for image processing after selection. 66,632 particles were autopicked via crYOLO and subjected to 2D classification in cryoSPARC. 57,798 selected particles were used for subsequent 3D classification in RELION-3.0.8, where the symmetrical *3up* map, described above, was used as initial reference. The best class comprising 22,055 particles (38%) represented an asymmetrical *1up/1up-out/1down* conformation with Sb#15 bound to each RBD. Several rounds of refinement and CTF refinement yielded a map of 4.0 Å resolution.

For the dataset of the S-6P/Sb#68 complex, in total 5,109 images were recorded, with 4,759 used for further image processing. 344,976 particles were autopicked via crYOLO and subjected to 2D classification in cryoSPARC. 192,942 selected particles were imported into RELION-3.0.8 and used for subsequent 3D classification, where the symmetrical *3up* map, described above, was used as initial reference. Two distinct classes of spike protein were found. One class (24,325 particles, 13%) revealed a state in which two RBDs adopt an *up* conformation with Sb#68 bound, whereby the density for the third RBD was poorly resolved representing an undefined state. Several rounds of refinement and CTF refinement yielded a map of 4.8 Å resolution. Two other classes, comprising 44,165 particles (23%) and 84,917 particles (44%), were identical. They show a *1up/2down* configuration without Sb#68 bound to any of the RBDs. Both classes were processed separately, whereby the class with over 80k particles yielded the best resolution of 3.3 Å and was used for further interpretation. A final non-uniform refinement in cryoSPARC further improved resolution down to 3.1 Å.

### Model building

Model building was carried out in COOT [72] using previously determined SARS-CoV-2 spike protein structures (PDB ID 7MY2, 6ZGG), Sb#68 crystal structure (PDB ID 7KLW) as reference and anti-GFP nanobody structure (PDB ID 3K1K) as a homology model for Sb#15. After each round of real-space refinement performed in Phenix [73], coordinates were manually inspected and edited in COOT and submitted to another refinement round in an iterative way.

## TABLES

**Table S1.**
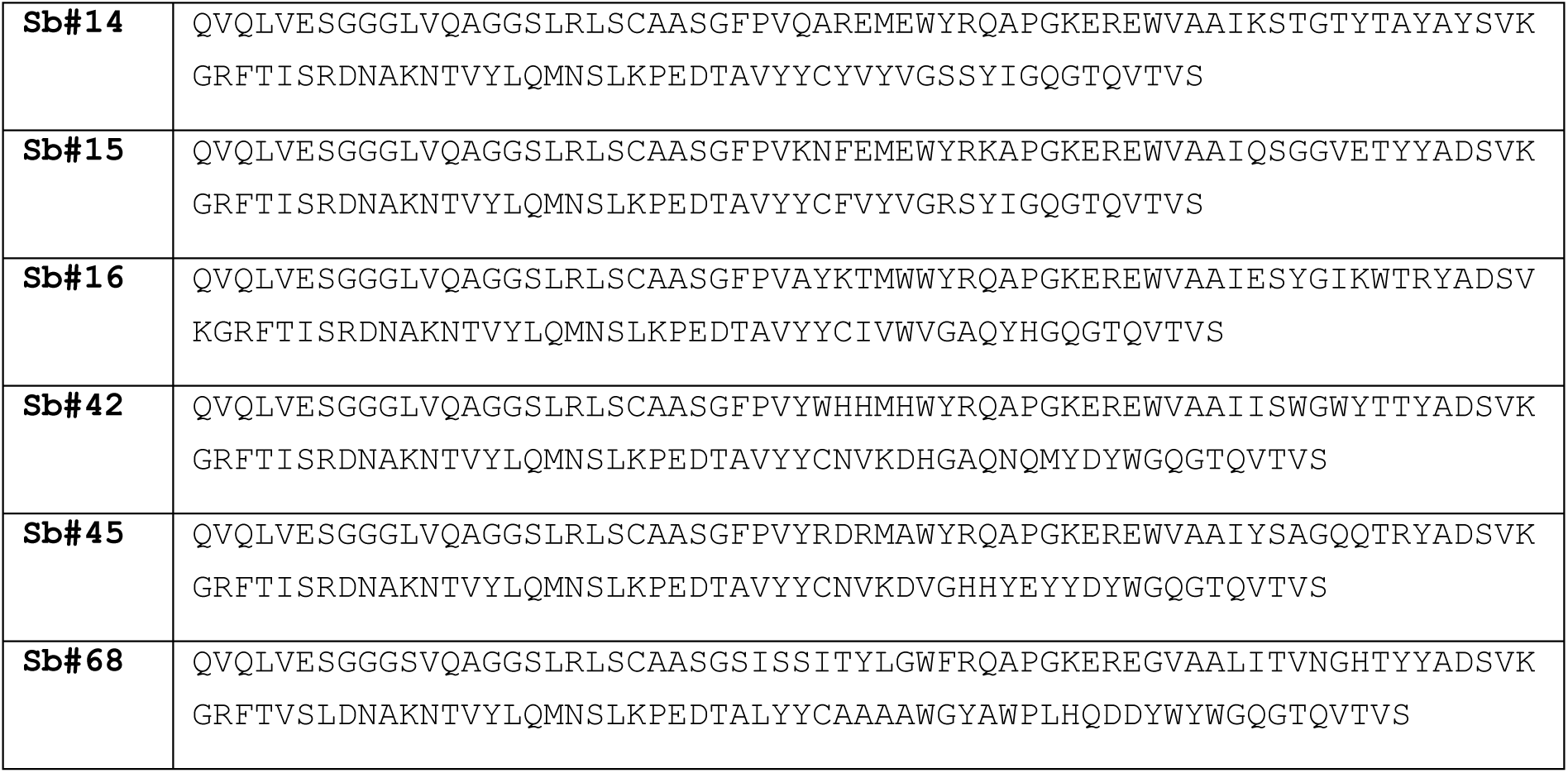
Sybody protein sequences

**Table S2.** Cryo-EM data collection, refinement and validation statistics

|  | S-2P + Sb#15 + Sb#68 |  | S-2P + Sb#15 | S-6P + Sb#68 |  |
| --- | --- | --- | --- | --- | --- |
|  | 3up<br>(EMD-12082,<br>PDB 7P77) | 1up/1up-out/1down<br>(EMD-12083, PDB<br>7P78) | 1up/1up-out/1down<br>(EMD-12084, PDB<br>7P79) | 2up/1flexible<br>(EMD-12085,<br>PDB 7P7A) | 1up/2down<br>(EMD-12086,<br>PDB 7P7B) |
| **Data collection and processing** |  |  |  |  |  |
| Magnification | 49,407 | 49,407 | 49,407 | 49,407 | 49,407 |
| Voltage (kV) | 200 | 200 | 200 | 200 | 200 |
| Electron exposure (e-/Å<sup>2</sup>) | 53 | 53 | 53 | 53 | 53 |
| Defocus range (μm) | -0.9 to -1.9 | -0.9 to -1.9 | -0.9 to -1.9 | -0.9 to -1.9 | -0.9 to -1.9 |
| Pixel size (Å) | 1.012 | 1.012 | 1.012 | 1.012 | 1.012 |
| Symmetry imposed | C3 | C1 | C1 | C1 | C1 |
| Initial particle images (no.) | 637,105 | 637,105 | 66,632 | 344,976 | 344,976 |
| Final particle images (no.) | 78,933 | 52,839 | 22,055 | 24,325 | 84,917 |
| Map resolution (Å) | 2.98 | 3.32 | 4.04 | 4.76 | 3.13 |
| FSC threshold | 0.143 |  |  |  |  |
| Map resolution range (Å) | 2.5 – 8.5 | 2.8 – 10.6 | 3.7-16.6 | 4.4-23.5 | 2.7-9.8 |
| **Refinement** |  |  |  |  |  |
| Initial model used | PDB 7MY2 | PDB 7MY2 | PDB 7MY2 | PDB 7MY2 | PDB 6ZGG |
| Model resolution (Å)<br>(0.5 FSC threshold) | 3.0 | 3.3 | 4.0 | 4.8 | 3.1 |
| Model resolution range (Å) | 15-3.0 | 15-3.3 | 17-4.0 | 24-4.8 | 15-3.1 |
| Map sharpening<br>B factor (Å<sup>2</sup>) | -99 | -82.7 | -80.4 | -118.3 | -92.2 |
| Model composition<br>Nonhydrogen atoms | 30582 | 28807 | 26907 | 25421 | 25096 |
| Protein residues | 3822 | 3625 | 3377 | 3159 | 3207 |
| Ligands | 66 | 56 | 56 | 65 | 0 |
| *B* factors (Å<sup>2</sup>) |  |  |  |  |  |
| Protein | 129.92 | 220.07 | 121.98 | 195.00 | 185.59 |
| Ligand | 114.69 | 157.41 | 138.65 | 217.68 | - |
| R.m.s. deviations |  |  |  |  |  |
| Bond lengths (Å) | 0.006 | 0.006 | 0.007 | 0.007 | 0.006 |
| Bond angles (°) | 1.059 | 1.078 | 1.220 | 1.190 | 1.005 |
| Validation |  |  |  |  |  |
| MolProbity score | 1.47 | 1.54 | 1.59 | 1.59 | 1.43 |
| Clashscore | 3.61 | 4.39 | 4.23 | 4.71 | 3.07 |
| Poor rotamers (%) | 0.00 | 0.00 | 0.00 | 0.00 | 0.04 |
| Ramachandran plot |  |  |  |  |  |
| Favored (%) | 95.46 | 95.39 | 94.39 | 95.09 | 95.24 |
| Allowed (%) | 4.54 | 4.58 | 5.61 | 4.95 | 4.73 |
| Disallowed (%) | 0.00 | 0.03 | 0.00 | 0.00 | 0.03 |

## DATA AVAILABILITY STATEMENT

The plasmids encoding for the six highest affinity binders are available through Addgene (Addgene #153522 - #153527). Purified Sb-Fc constructs can be commercially obtained from Absolute Antibody. The three-dimensional cryo-EM density maps have been deposited in the Electron Microscopy Data Bank under accession numbers EMD-12082 (SARS-CoV-2 spike protein in complex with Sb#15 and Sb#68 in a *3up* conformation), EMD-12083 (SARS-CoV-2 spike protein in complex with Sb#15 and Sb#68 in a *1up/1up-out/1down* conformation), EMD-12084 (SARS-CoV-2 spike protein in complex with Sb#15 in a *1up/1up-out/1down* conformation), EMD-12085 (SARS-CoV-2 spike protein in complex with Sb#68 in a *2up/1flexible* conformation) and EMD-12086 (SARS-CoV-2 spike protein in complex with Sb#68 in a 1up/2down conformation) and include the cryo-EM maps, both half-maps, the unmasked and unsharpened refined maps and the mask used for final FSC calculation. Coordinates of the models have been deposited in the Protein Data Bank. The accession numbers are 7P77, 7P78, 7P79, 7P7A, 7P7B, respectively.

## AUTHOR CONTRIBUTIONS

JDW, PP and MAS conceived the project. JDW and IZ cloned and purified target proteins. CAJH performed the sybody selection, made the bispecific constructs and purified sybodies. JDW and CAJH performed ELISA and GCI experiments. AAG and JR acquired and processed cryo-EM data under the supervision of CP and DJS using proteins prepared by JDW and CAJH. MS and MW performed neutralization assays under the supervision of PP using pseudotyped VSV and sybodies produced by CAJH and JDW. YR performed the neutralization experiments with live SARS-CoV-2 under the supervision of GZ. JCE prepared GFP-purification resin. PE, MS and IZ were involved in planning the sybody selection strategy. All authors discussed and interpreted data. JDW, CAJH, AAG, MS and MAS prepared figures. JDW, CP, PP and MAS wrote the paper. CAJH, AAG, GZ and DJS made major contributions in the course of paper editing.

## ACKNOWLEDGEMENTS

We thank Rony Nehmé and André Heuer (Creoptix AG, Wädeswil, Switzerland) for the acquisition, fitting and interpretation of a first set of GCI measurements using the WAVEsystem. We thank Florence Projer, David Hacker and Kelvin Lau (Protein Production and Structure Core Facility, EPFL, Switzerland) for the production of the pre-fusion spike protein. We are grateful to Jason McLellan (The University of Texas at Austin, U.S.) for having provided the pre-fusion-stabilized soluble spike expression vectors for S-2P and S-6P. We thank Michael Fiebig (Absolute Antibody) for providing us with purified Sb-Fc. We thank Raimund Dutzler and Marta Sawicka (University of Zurich) for freezing cryo-EM grids. Michiel Punter (University of Groningen) is acknowledged for IT help. We are also grateful to the Next Generation Sequencing Platform, University of Bern for their contribution to this study.

---

**Figure S1.**
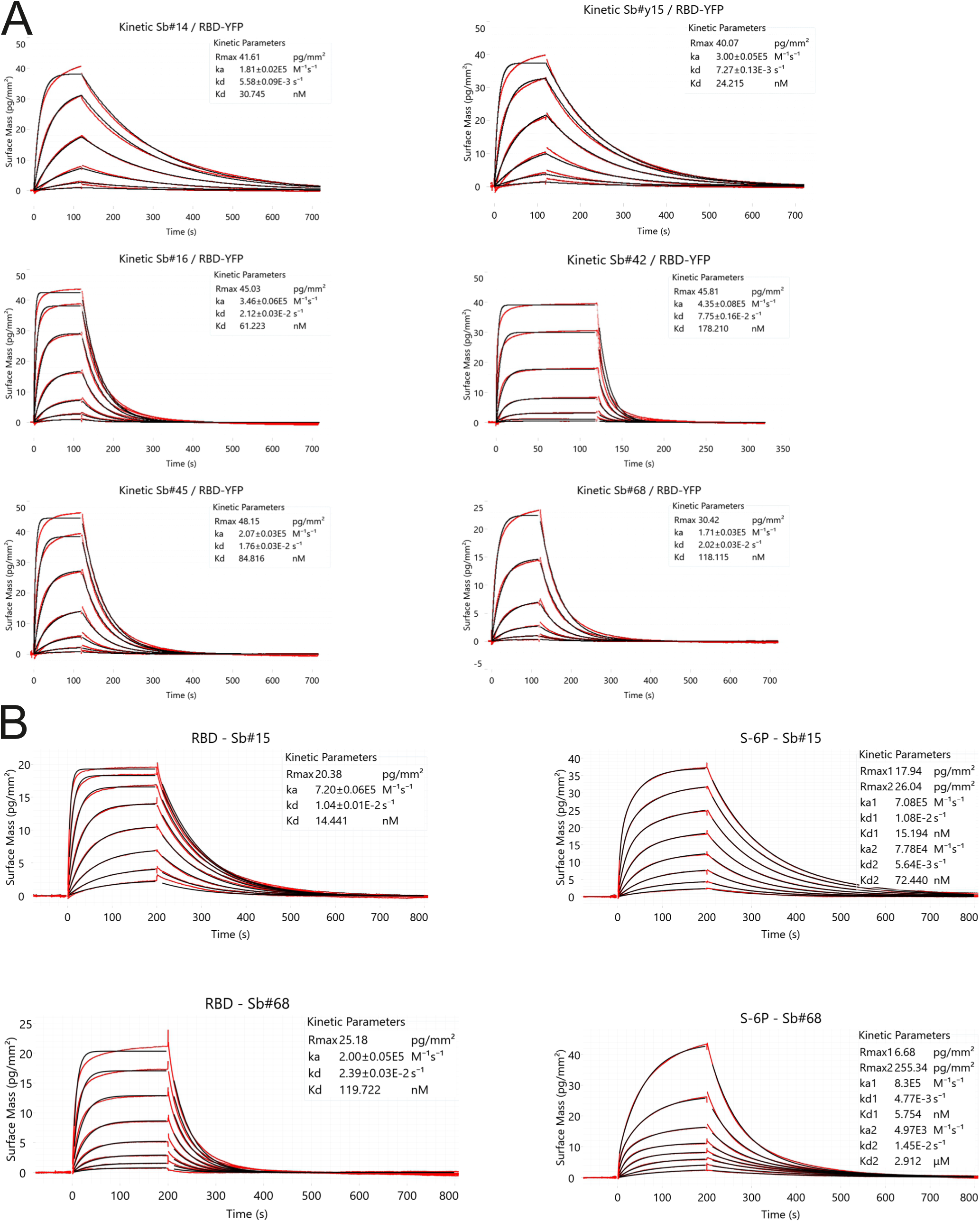
Kinetic characterization of sybodies by GCI. (**A**) RBD-vYFP and ECD were immobilized as indicated and the six top sybodies were injected at increasing concentrations ranging from 1.37 nM to 1 μM. Data were fitted using a Langmuir 1:1 model. (**B**) In depth affinity characterization of Sb#15 and Sb#68. RBD-vYFP and S-6P were immobilized as indicated and Sb#15 and Sb#68 were injected at concentrations ranging from 1.95 nM to 250 nM for Sb#15 and 3.9 nM to 500 nM for Sb#68. For RBD, data were fitted using a Langmuir 1:1 model. For S-6P, the data were fitted with the heterogeneous ligand model, because the 1:1 model was clearly not appropriate to describe the experimental data. Corresponding data for S-2P is shown in main Fig. 1C.

**Figure S2.**
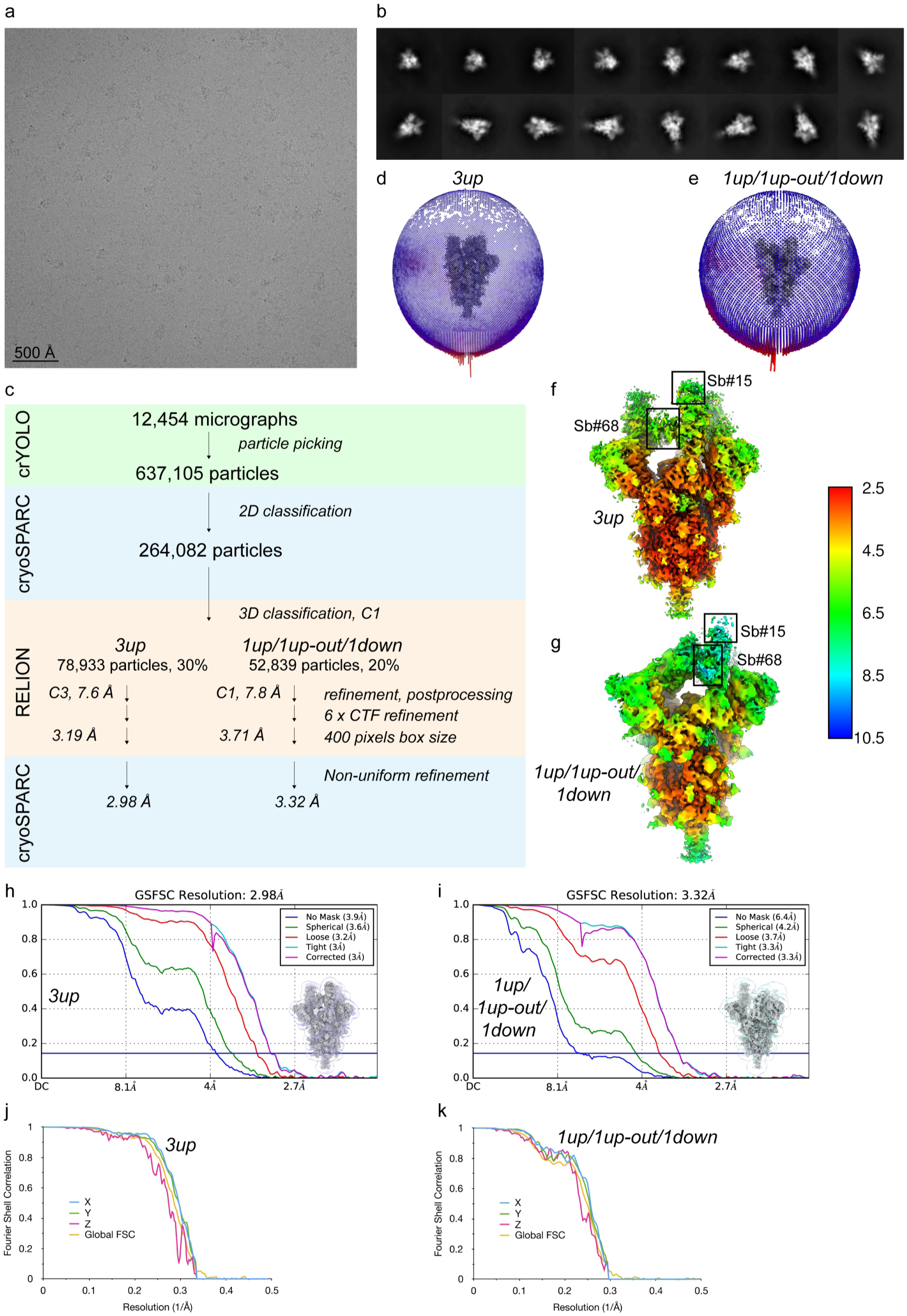
Cryo-EM reconstruction of S-2P in complex with Sb#15 and Sb#68. Representative cryo-EM image (**A**) and 2D-class averages (**B**) of vitrified S-2P in the presence of both Sb#15 and Sb#68. (**C**) Image processing work flow. Angular particle distribution plot (**D** and **E**), final reconstructed map colored by local resolution, as estimated in cryoSPARC (**F** and **G**), FSC plot (**H** and **I**) and anisotropy plot used for resolution estimation (**J** and **K**) for the final 3*up* and 1*up*/1*up-out*/1*down* RBD reconstruction, respectively. (**H** and **I**) The line indicates the FSC thresholds used for FSC of 0.143, and the mask used for FSC calculation overlaid on the map is shown as thumbnail. (**J** and **K**) The global FSC curve is represented in yellow, while the directional FSCs along the x, y and z axis are displayed in blue, green and red, respectively.

**Figure S3.**
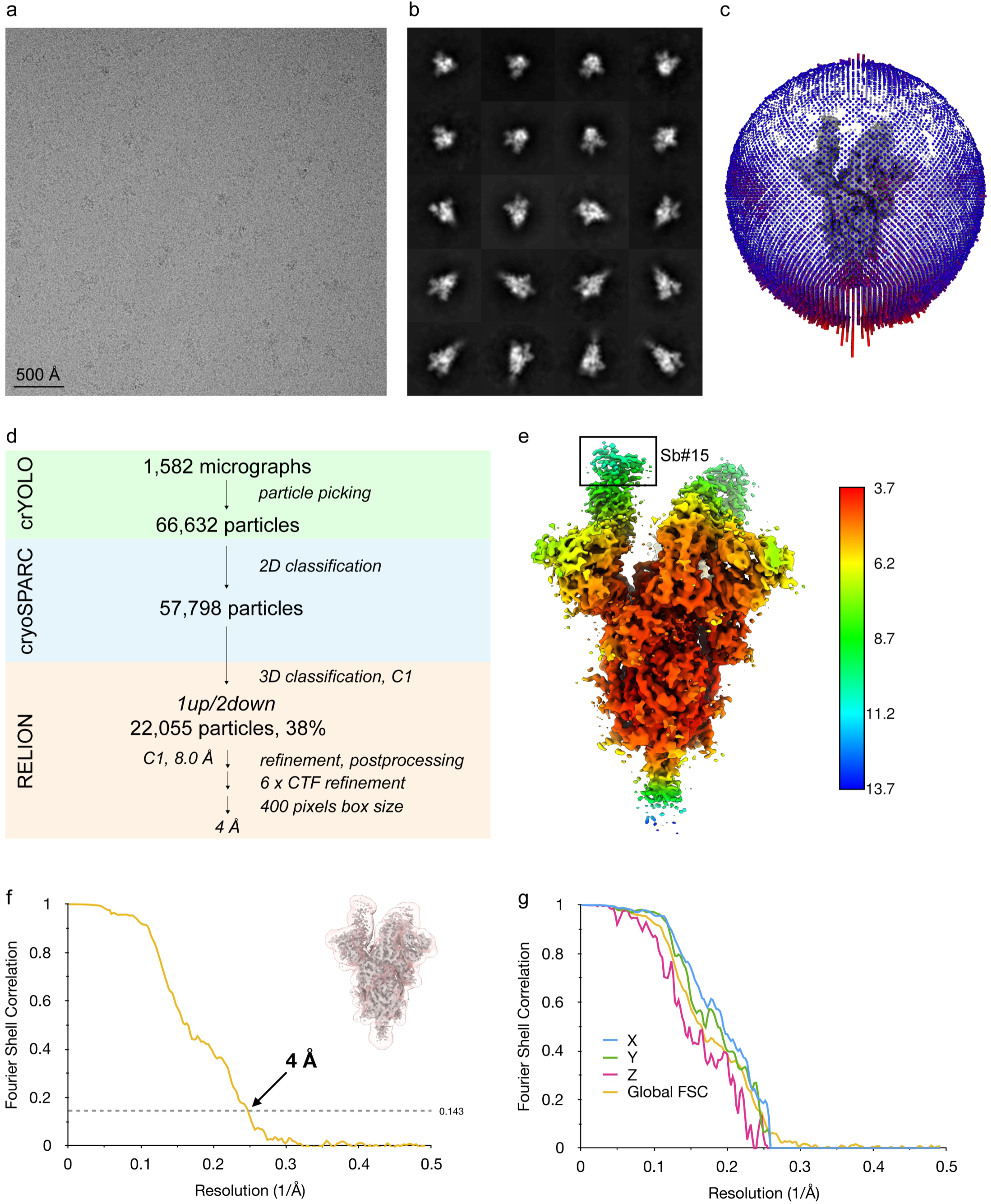
Cryo-EM reconstruction of S-2P in presence of Sb#15. Representative cryo-EM image (**A**) and 2D-class averages (**B**) of vitrified S-2P in the presence of Sb#15. (**C**) Angular distribution plot of particles included in the final map. (**D**) Image processing work flow. Final reconstructed map colored by local resolution, as estimated in RELION (**E**), FSC plot (**F**) and anisotropy plot used for resolution estimation (**G**) for the final 1*up*/2*down* RBD reconstruction. (**F**) The line indicates the FSC thresholds used for FSC of 0.143, and the mask used for FSC calculation overlaid on the map is shown as thumbnail. (**G**) The global FSC curve is represented in yellow, while the directional FSCs along the x, y and z axis are displayed in blue, green and red, respectively.

**Figure S4.**
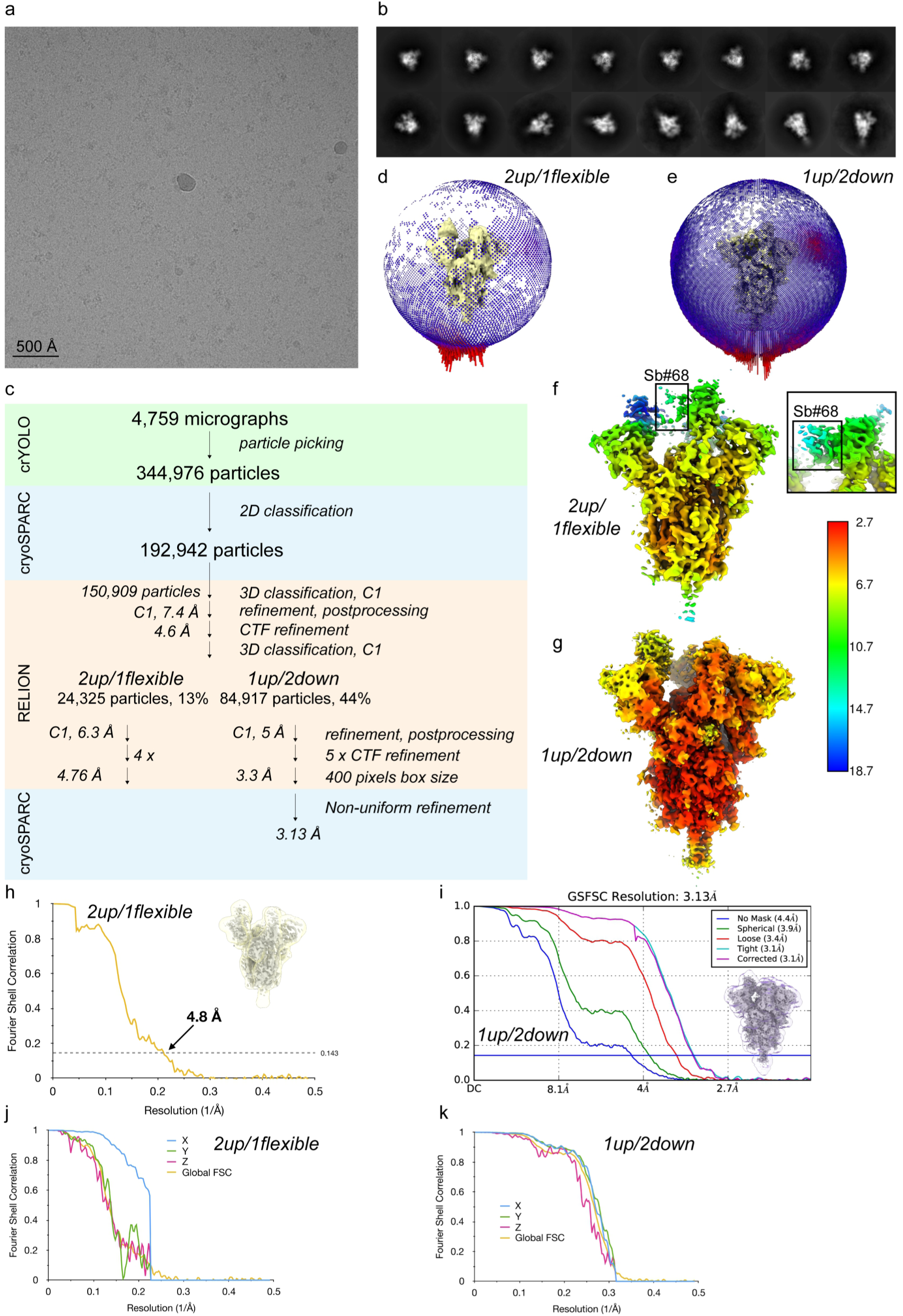
Cryo-EM reconstruction of S-6P in presence of Sb#68. Representative cryo-EM image (**A**) and 2D-class averages (**B**) of vitrified S-6P in the presence of Sb#68. (**C**) Image processing work flow. Angular particle distribution plot (**D** and **E**), final reconstructed map colored by local resolution, as estimated in cryoSPARC and RELION (**F** and **G**), FSC plot (**H** and **I**) and anisotropy plot used for resolution estimation (**J** and **K**) for the final 2*up*/1*flexible* and 1*up*/2*down* RBD reconstruction, respectively. (**H** and **I**) The line indicates the FSC thresholds used for FSC of 0.143, and the mask used for FSC calculation overlaid on the map is shown as thumbnail. (**J** and **K**) The global FSC curve is represented in yellow, while the directional FSCs along the x, y and z axis are displayed in blue, green and red, respectively.

**Figure S5.**
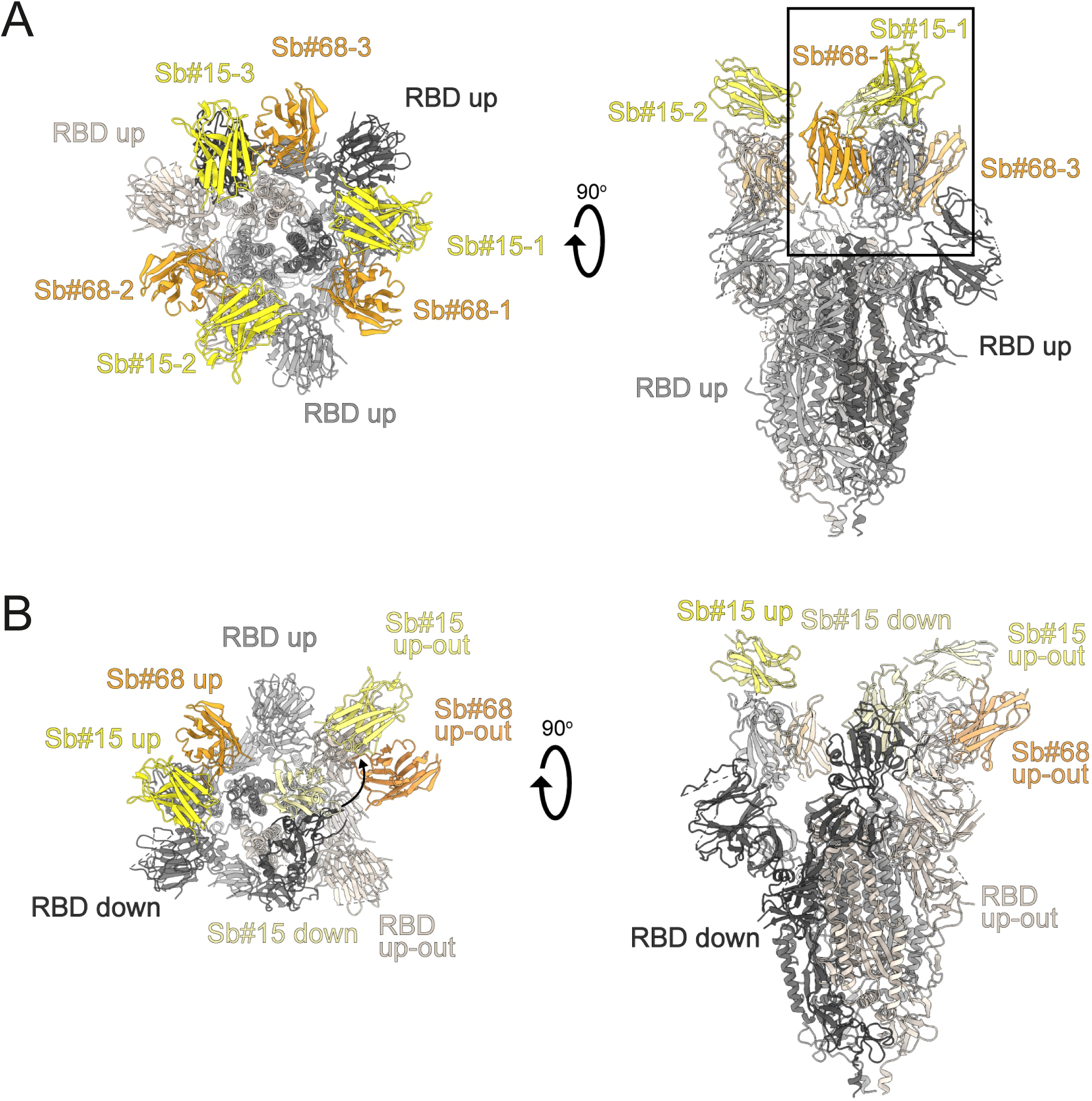
Structures of S-2P spike in complex with both Sb#15 and Sb#68. (**A**) Structure of S-2P with both Sb#15 and Sb#68 bound to each RBD adopting a symmetrical 3*up* conformation. Based on the cryo-EM map shown in main Fig. 5A, a model shown as ribbon was built using pre-existing structures (PDB ID:6X2B for S-2P; PDB ID:3K1K for Sb#15; PDB ID:5M13 for Sb#68). (**B**) Structure of S-2P with the three RBDs adopting an asymmetrical 1*up*/1*up-out*/1*down* conformation. Based on the cryo-EM map shown in Fig. 5B, a model shown as ribbon was built using pre-existing structures (PDB ID:6X2B for S-2P; PDB ID:3K1K for Sb#15; PDB ID:5M13 for Sb#68). The *up-out* state is pushed outward by the adjacent RBD in a down state with bound Sb#15 (arrow). Spike protein is shown in shades of grey, Sb#15 in yellow and Sb#68 in orange.

**Figure S6.**
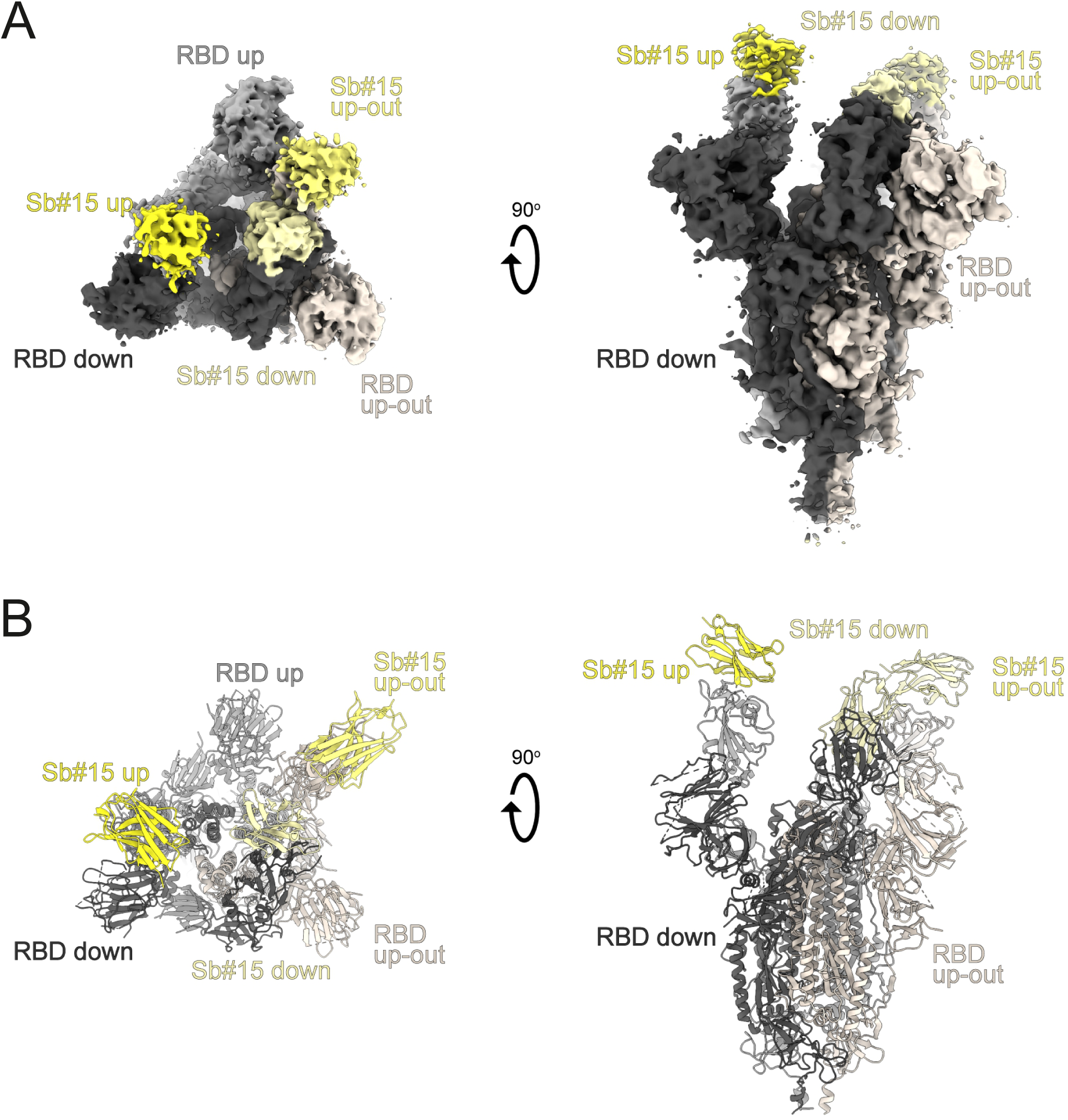
Structural analysis of the S-2P/Sb#15 complex. (**A**) Cryo-EM map of S-2P with Sb#15 bound to each RBD adopting an asymmetrical 1*up*/1*up-out*/1*down* conformation. (**B**) The corresponding model built using pre-existing structures (PDB ID:6X2B for S-2P; PDB ID:3K1K for Sb#15) is shown as ribbon. Final map blurred to a B factor of -30 Å was used for better clarity of the less resolved RBDs and sybodies. Spike protein is shown in shades of grey and Sb#15 in yellow.

**Figure S7.**
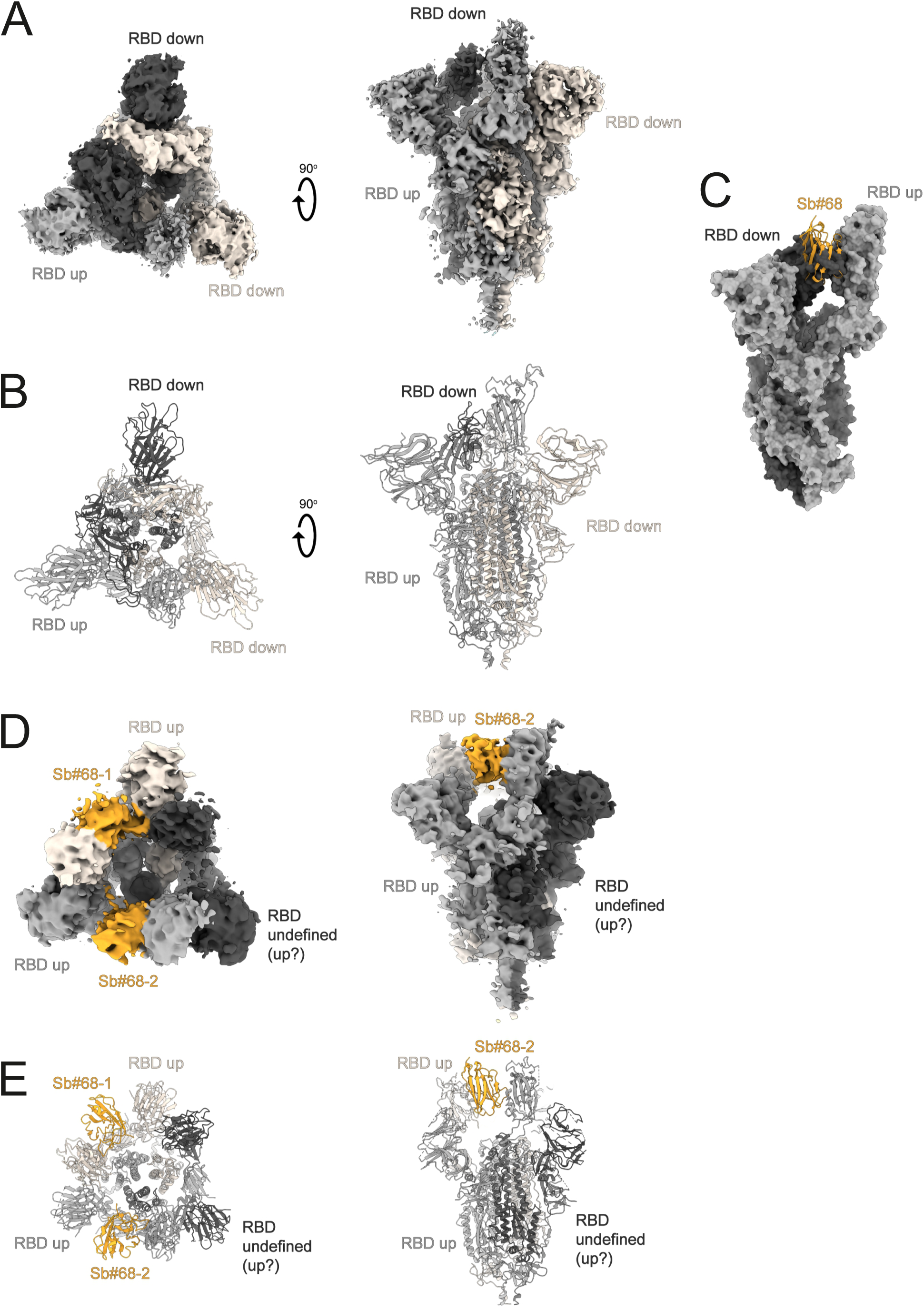
Structural analysis of S-2P/Sb#68 complex. (**A**) Cryo-EM map of S-6P, with an 1*up*/2*down* RBD conformation. (**B**) The corresponding model is shown as ribbon (PDB ID:6ZGG for S-6P). No densities for Sb#68 were observed. (**C**) Sb#68 cannot bind to *up* RBD if the neighbouring RBD exhibits a *down* conformation due to steric clashing. (**D**) Cryo-EM map of S-6P, with two Sb#68 bound to *up* RBDs of the spike featuring a 2*up*/1*flexible* conformation. (**E**) The corresponding model built on pre-existing structures (PDB ID:6X2B for S-6P; PDB ID:5M13 for Sb#68) is shown as ribbon. Final maps blurred to a B factor of -30 Å were used for better clarity of the less resolved RBDs and sybodies. Spike protein is shown in shades of grey and Sb#68 in orange.

**Figure S8.**
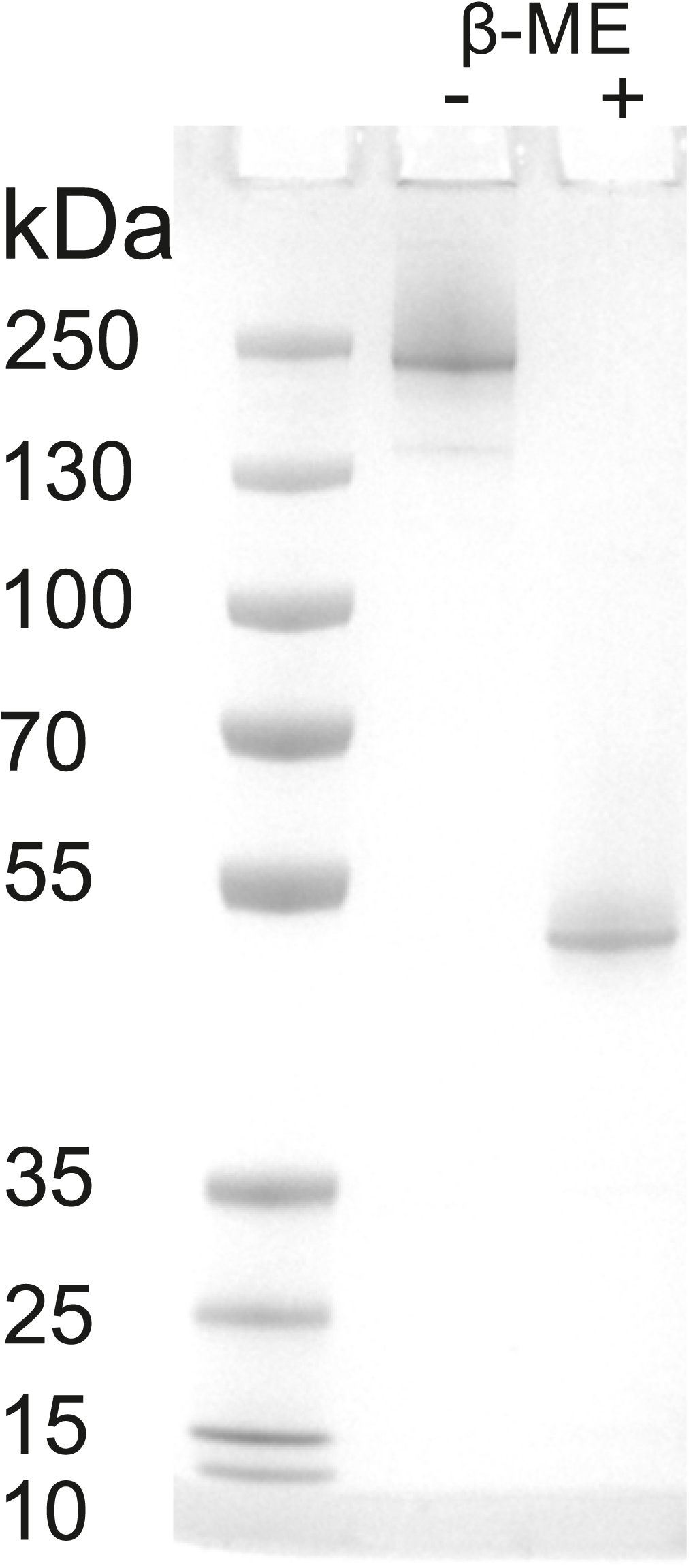
SDS-PAGE analysis of Tripod-GS4r. Purified Tripod-GS4r was loaded on a SDS-PAGE gel with and without incubation of β-mercaptoethanol (β-ME) and stained using coomassie blue.

## Notes

### Competing Interest Statement

Iwan Zimmermann, Pascal Egloff and Markus A. Seeger are founders and shareholders of Linkster Therapeutics AG.

### Summary of Updates

In relation to the initial version, this version contains escape mutant data and data involving variants of concern.

